# Dissecting Gene Regulatory Networks Governing Human Cortical Cell Fate

**DOI:** 10.1101/2025.09.23.678137

**Authors:** Jingwen W. Ding, Chang N. Kim, Megan S. Ostrowski, Yashodara Abeykoon, Bryan J. Pavlovic, Jenelle L. Wallace, Tomasz J. Nowakowski, Alex A. Pollen

## Abstract

Human cortical neurogenesis involves conserved and specialized developmental processes during a restricted window of prenatal development. Radial glia (RG) neural stem cells shape cortical cell diversity by giving rise to excitatory neurons, oligodendrocytes, and astrocytes, as well as olfactory bulb interneurons (INs) and a recently characterized population of cortical INs^1,2^. Complex genetic programs orchestrated by transcription factor (TF) circuits govern the balance between self-renewal and differentiation, and between different cell fates^3–8^. Despite progress in measuring gene regulatory network activity during human cortical development^9–12^, functional studies are required to evaluate the roles of TFs and effector genes in human RG lineage progression. Here we establish a human primary culture system that allows sensitive discrimination of cell fate dynamics and apply single cell clustered regularly interspaced short palindromic repeats interference (CRISPRi) screening^13,14^ to examine the transcriptional and cell fate consequences of 44 TFs active during cortical neurogenesis. We identified multiple TFs, with novel roles in cortical neurogenesis, including *ZNF219,* previously uncharacterized, that represses neural differentiation and *NR2E1* and *ARX* that have opposing roles in regulating RG lineage plasticity and progression across developmental stages. We also uncovered convergent effector genes downstream of multiple TFs enriched in neurodevelopmental and neuropsychiatric disorders and observed conserved mechanisms of RG lineage plasticity across primates. We further uncovered a postmitotic role for *ARX* in safeguarding IN subtype specification through repressing *LMO1*. Our study provides a framework for dissecting regulatory networks driving cell fate consequences during human neurogenesis.

## Main

Human radial glia (RG) evolved an increased proliferative capacity, altered cell fate potential, and protracted period of maturation, supporting the increased number and complexity of daughter cells^2,15,16^. Gene regulatory networks governing RG self-renewal, differentiation, and maturation have long been implicated in cortical expansion^17,18^. RG sequentially produce distinct subtypes of neurons followed by glial cell types^19,20^. Although cortical inhibitory neurons (INs) are mainly generated in the ganglionic eminences (GE) and migrate to the cortex^21–23^, recent lineage tracing studies^1,24–26^ and developmental cell atlases^27^ have demonstrated that human RG also produce cortical-like INs at late stages of neurogenesis that share transcriptional signatures with caudal and lateral GE (CGE and LGE)-derived INs. However, the role of TFs in regulating RG differentiation decisions, lineage plasticity to produce INs, and maturation remains largely unexplored.

Genetic perturbations combined with measurements of single cell gene expression provide a powerful approach, termed Perturb-seq, for high-throughput dissection of gene function^13,14^. Cas9-based CRISPR loss-of-function perturbation approaches have recently been applied to study gene regulatory networks influencing cortical development using induced pluripotent stem cell (iPSC)-derived organoid^28–30^ and mouse^31,32^ models. However, Cas9-induced double-stranded DNA breaks can cause cytotoxicity, influencing proliferation and differentiation decisions^33,34^, while acquired genetic and epigenetic variation in source iPSCs^35,36^, and patterning biases^37^ and cell stress^38^ during differentiation can influence cell type fidelity and fate specification in organoid models. CRISPR interference (CRISPRi) screens using dCas9-KRAB enable efficient and uniform repression of target genes with limited cellular toxicity^39,40^, while primary cortical culture^41,42^ captures physiological aspects of human cortical neurogenesis, supporting sensitive detection of cell fate choices.

Here we established a primary cell model system that recapitulates *in vivo* differentiation dynamics and performed Perturb-seq to measure the transcriptional and cell fate consequences of repressing 44 TFs that are robustly expressed in the human cortical RG lineage. Extending human primary cell culture approaches^1,42,43^, we targeted TFs in a homogeneous RG population and then removed growth factors to permit cortical neurogenesis, cell fate choice, and early subtype specification. Our screening revealed the role of *ZNF219*, not previously described in cortical development, in repressing neuronal differentiation, opposing roles for *NR2E1* and *ARX* in regulating the balance of human excitatory versus inhibitory neurogenesis, and the role of *ARX* in safeguarding IN subtype specification through transcriptional repression of downstream transcription cofactor *LMO1*. Intersecting dysregulated genes under different perturbations revealed candidate hub effector genes downstream of multiple TFs enriched for roles in neurodevelopmental and neuropsychiatric disorders. Coupling CRISPRi screening with barcoded lineage tracing demonstrated the potential to engineer lineage plasticity and developmental tempo of individual RG through TF perturbation. Together with parallel screening in rhesus macaque, our data illuminate conserved mechanisms governing cortical RG lineage progression across primates.

## A primary model that enables systematic TF repression during human cortical neurogenesis

We designed a primary cell model of neurogenesis and lineage progression to evaluate the impacts of TF repression on cell fate choice during human cortical development. To direct gene targeting to RG at the start of lineage progression, we first enriched for RG isolated from primary human tissue samples by adding epidermal growth factor (EGF) and fibroblast growth factor 2 (FGF2) for five days prior to infecting with an all-in-one CRISPRi lentivirus (Fig. 1a), and we further expanded RG for one week allowing for target gene knockdown prior to differentiation^39^ (Extended Data Fig. 1a-c). We then replaced growth factors with brain derived neurotrophic factor (BDNF) to support spontaneous differentiation of perturbed RG. Consistent with recent studies^1,42,43^, this model recapitulates *in vivo* RG lineage progression and generation of ENs and INs (Fig. 1a and Extended Data Fig. 1a-b), enabling detection of perturbations that affect differentiation dynamics and cell fate choice. Analysis of *PAX6* and *EOMES* recapitulated the effects of these TFs described in other model systems in promoting excitatory neurogenesis, highlighting the potential of our system to uncover additional regulators^44–47^ (Extended Data Fig. 1d).

**Fig. 1:**
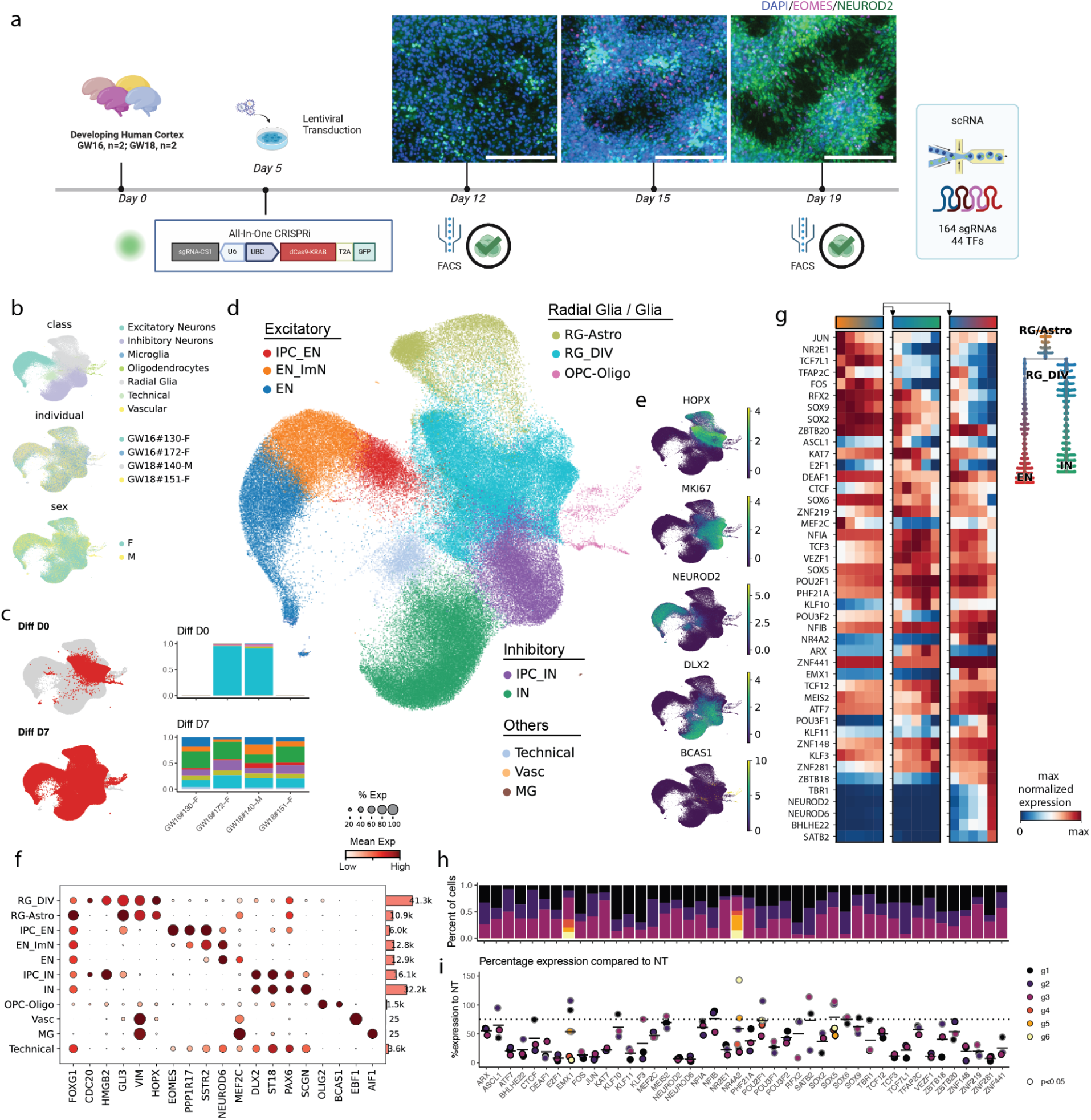
Single cell Perturb-seq on 44 TFs in primary human neuronal differentiation. a. Experimental design for high-throughput perturbation of 44 TFs in human primary cortical progenitors from 4 individuals, with immunocytochemical labeling of *EOMES* and *NEUROD2* in *in vitro* culture of human cortical RG before (D0) and after induced differentiation (D3, D7). b. UMAPs of cells collected on D0 (21,151 cells, N = 2 individuals) and D7 (116,166 cells, N = 4 individuals), colored by cell class, individual and sex. c. UMAPs with cells from different time points highlighted. d. UMAP colored by supervised cell type, with stacked barplots (bottom left) showing distributions of cell types at each stage and in each individual with the same colors, split by timepoints. e. UMAPs colored by the expression of *HOPX, MKI67, NEUROD2, DLX2* and *BCAS1*. f. Dotplot of the expression of cell type markers for the assigned clusters (left), with a barplot showing numbers of cells detected in the cluster (right). The size of each dot denotes the fractions of cells in the group where the gene is expressed and the color denotes mean gene expression in the group. g. Heatmap of normalized expression of 44 TFs targeted in this study along pseudotime with branches for IN and EN (left), with a tree plot showing cell type distribution along pseudotime with lineage branches (right). Pseudotime was calculated using NT cells on D7. h. Stacked barplot of detected sgRNAs for each TF target on D7, colored by individual sgRNA. i. Dotplot for percentage expression of target gene expression compared to NT cells on D7, filled by individual sgRNA, with borders highlighting significance. log_2_FC were calculated in each cell class and the lowest value for each sgRNA was used for visualization and filtering. sgRNAs that reduced expression by less than 25% compared to NT were considered inactive and removed from downstream analyses and remaining active sgRNAs had a median expression of 28% (72% KD). Scale bars: 200 μm GW, gestational week; sgRNA, single guide RNA; TF, transcription factor; FACS, fluorescence-activated cell sorting; RG, radial glia; Astro, astrocytes; EN, excitatory neuron; IN, inhibitory neuron; IPC, intermediate progenitor cell; ImN, immature neuron; DIV, dividing; OPC, Oligodendrocyte progenitor cell; MG, microglia; Vasc, vascular; NT, non-targeting; log_2_FC, log_2_ fold change

To systematically identify regulators of human cortical neurogenesis, we first prioritized TFs using single cell RNA sequencing (scRNA-seq) and single cell assay for transposase-accessible chromatin sequencing (scATAC-seq) multiome data. We selected 44 TFs based on robust expression, motif accessibility, gene regulatory network size, target gene expression and predicted transcriptional consequences in the RG lineage (Methods, Supplementary Table 1). We then adapted an all-in-one CRISPRi vector co-expressing green fluorescent protein (GFP)^48^ by inserting a capture sequence in the single guide RNA (sgRNA) scaffold, enabling direct capture of sgRNAs during scRNA-seq using a 10X Genomics platform^49^, thereby supporting screening in primary cell culture systems. We synthesized and cloned a library containing 164 sgRNAs, targeting the 44 TFs, and included 20 non-targeting control (NT) sgRNAs. For each TF, we targeted active promoters with accessible chromatin during human cortical neurogenesis^9^ using 3 sgRNAs per promoter (Supplementary Table 2)^50,51^. We derived primary human cultures from cryopreserved cortical tissue of four individuals at stages of peak neurogenesis from gestational weeks (GW)16-18^41^. We delivered the CRISPRi library by lentivirus into primary RG, targeting less than 30% infection rate to generate libraries with a majority of singly infected cells (Extended Data Fig. 1a, Supplementary Table 3), removed growth factors, and confirmed efficient gene repression prior to and throughout differentiation (Extended Data Fig. 1c,e).

scRNA-seq confirmed a 95% population of cycling RG on day 0 of differentiation (D0), marked by co-expression of RG marker *HOPX* and proliferation marker *MKI67* (Fig. 1b-e). By D7, three principal cortical cell class trajectories emerged - ENs, INs and oligodendrocytes marked by *NEUROD2*, *DLX2, BCAS1*, respectively (Fig. 1c-e), in addition to a continuum from RG to astrocytes marked by *AQP4* (Extended Data Fig. 1f). Integrating both experimental timepoints highlighted homogeneity of D0 populations with minimal spontaneous differentiation (Fig. 1c, Extended Data Fig. 1g), as well as comparable representation of cells derived from independent technical and biological replicates across cell types with minimal batch effects (Fig. 1b and Extended Data Fig. 1h).

Pearson correlation and reference mapping to a developmental cortical cell atlas^12^ confirmed the recapitulation of *in vivo*-like gene expression, cell types, states, differentiation dynamics, and cell fate choice in the primary culture differentiation system (Extended Data Fig. 1i,j). The majority of neurons exhibited immature states (Extended Data Fig. 1j), consistent with recent generation from RG after differentiation. These differentiation dynamics were also recovered by RNA velocity and pseudotime analysis (Fig. 1g and Extended Data Fig. 1l,m). Notably, comparison with iPSC based models revealed a near 2-fold increase in the Pearson correlation of marker gene expression to *in vivo* cell types in the primary 2D system, supporting improved fidelity to normal development (Extended Data Fig. 1i,k). In addition, primary culture in 2D and organotypic slice both showed reduced transcriptional signatures of cellular stress across major dimensions, including glycolysis, endoplasmic reticulum (ER) stress, oxidative stress, and apoptosis^52,53^ (Extended Data Fig. 1n), supporting the physiological relevance of the primary cell models.

We assigned sgRNAs to 118,456 cells (Methods) across timepoints, with a mean of 200 singly infected cells per sgRNA on differentiation D7 (Fig. 1h and Extended Data Fig. 1o). Knockdown (KD) efficiency was calculated using DEseq2^54^ in each cell class and 18 sgRNAs with less than 25% KD (log_2_ fold change > -0.4) were removed from downstream analyses (Fig. 1i and Supplementary Table 4). Remaining active sgRNAs exhibited a median KD efficiency of 72%, with comparable transcriptional responses to independent sgRNAs targeting the same promoter (Fig. 1i and Extended Data Fig. 2). For further analysis, we collapsed all active sgRNAs sharing the same target TF, which yielded a median KD efficiency of 80% and a mean of 600 cells per gene (Extended Data Fig. 1o,p).

## Convergent transcriptional consequences reveal enrichment of disease related effector genes

We next examined the consequences of TF repression on gene expression and cell type composition by differentiation D7 (Fig. 2a and Supplementary Table 5). Gene expression changes were correlated between individual sgRNAs and genes (average Pearson R = 0.88) and cell type abundance was preserved upon downsampling of cell number (average Pearson R = 0.94), supporting the power of the screen to detect changes in both modalities (Extended Data Fig. 3a,b).

**Fig. 2:**
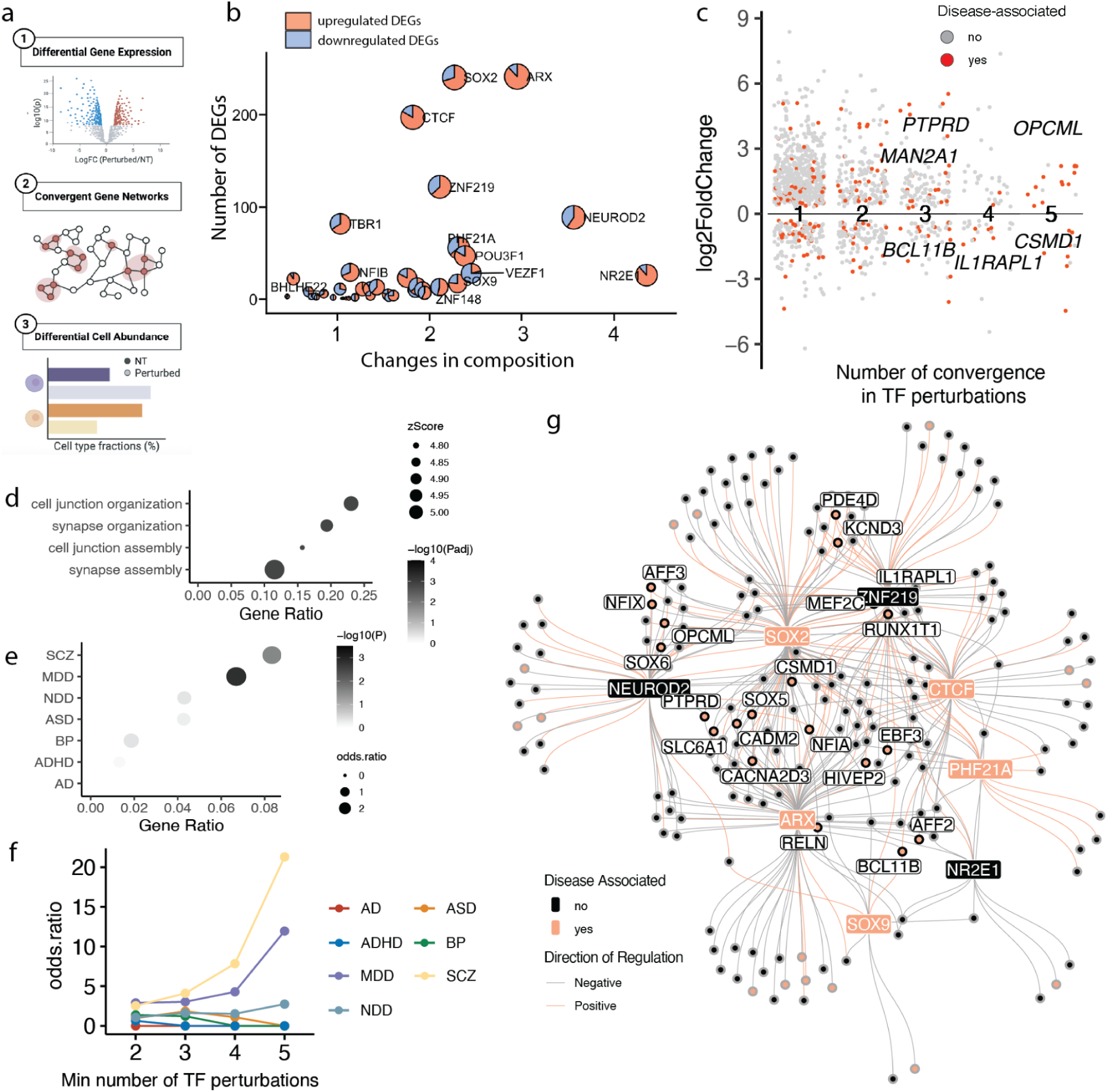
Convergent transcriptional changes reveal neuropsychiatric disorder -enriched hub effector genes. a. Schematics to show analytical workflow applied to the D7 Perturb-seq data obtained from Fig. 1. b. Scatterplot for summed absolute values of estimated coefficient in cell abundance (X axis) and number of DEGs (Y axis) across 7 cell types in the EN and IN lineage shown in Fig. 2a for each TF perturbation. Dot size reflects perturbations with the strongest phenotypes calculated by (summed absolute coefficients * total number of DEGs), and each dot includes a pie chart colored by the fraction of up or down regulated DEGs. c. Volcano plot for DEGs shared between different numbers of perturbations. DEGs with an absolute log2FC change more than 0.2 were shown. DEGs that overlap with the 7 sets of disease associated genes were colored as red and DEGs associated with SCZ are labeled. d. Dotplot for GO enrichment in biological processes of convergent DEGs that were detected in at least 2 TF perturbations using non-convergent DEGs as background. Dot size denotes Z score and color denotes -log_10_(adjusted p value). e. Dotplot for enrichment of 7 sets of disease associated genes (Methods) in convergent DEGs identified by Fisher’s exact test compared to non-convergent DEGs. Dot size denotes odds ratio and color denotes -log_10_(p value). f. Line plot for odds ratio of disease associated gene enrichment in DEGs identified in greater than or equal to 2, 3, 4, 5 perturbations. Color indicates different sets of disease genes. g. Regulatory network plot to show 8 prioritized TFs (*NR2E1, ARX, SOX2, ZNF219, NEUROD2*, *PHF21A, SOX9* and *CTCF*) and connected convergent DEGs. Color of the edges indicate the direction of regulation (pink suggests positive regulation and that the DEG was downregulated upon TF perturbation). Pink nodes/labels represent disease-associated genes/TFs. FC,fold change; DEGs, differentially expressed genes; NT, non-targeting; SCZ, schizophrenia; MDD, major depressive disorder; ASD, autism disorder; NDD, neurodevelopmental disorder; ADHD, attention-deficit/hyperactivity disorder; BP, bipolar disorder; AD, Alzheimer’s disease

We observed a positive correlation between the effects of individual TFs on cell type composition and on gene expression across cell types (Fig. 2b) and within cell classes (Extended Data Fig. 3c). Comparing the extent of changes in both dimensions prioritized TFs whose depletion caused the strongest phenotypes: *NR2E1*, *ARX*, *ZNF219*, *SOX2*, *SOX9*, *CTCF*, *NEUROD2* and *PHF21A* (Fig. 2b). These TFs also showed the strongest impact on other molecular phenotypes, including Euclidean distance and energy distance^55^ to NT, which compare the average expression between groups and the distance between and within groups, respectively, as well as maximum composition change among all cell types in the RG lineage (Extended Data Fig. 3d). Notably, the TFs with strong cellular and transcriptional phenotypes have previously been implicated in neurological disorders: *ARX* in X-linked lissencephaly^56^, epilepsy^57,58^ and intellectual disability^59,60^, and autism spectrum disorder (ASD)^57^; *NR2E1* in schizophrenia (SCZ)^61^; *SOX2* in intellectual disability and epilepsy^62,63^; *CTCF* in intellectual disability with microcephaly^64,65^; *NEUROD2* in intellectual disability^66^ and early infantile epileptic encephalopathy^67^; *PHF21A* in intellectual disability with epilepsy and ASD^68^ and *ZNF219* in a case of low IQ ASD^69^. While we cannot rule out that additional TFs may impact differentiation or maturation phenotypes not captured in this assay due to incomplete knockdown, redundancy, or stage-specific roles, these results highlight the functional importance and disease relevance of TFs prioritized by screening.

At the gene expression level, the predominance of up-regulated DEGs following *NR2E1* (88%) and *ARX* (85%) KD was consistent with their role as transcriptional repressors^70,71^ (Fig. 2b). Approximately 25% of DEGs were affected by the perturbation of more than one TF (Fig. 2c). These convergent DEGs showed enrichment for cell adhesion and synaptic development related terms, suggesting that effector genes regulating neurogenesis, migration and maturation are highly interconnected across TF regulatory networks (Fig. 2d). Moreover, intersecting these convergent DEGs with 7 sets of neurological and neurodevelopmental disorder-related genes revealed significant overlaps with SCZ and major depressive disorder (MDD)-associated genes, in comparison to non-convergent DEGs (Fisher’s exact test, p < 0.05) (Fig. 2e, f). This overlap includes *PTPRD*, modulated by *ARX*, *SOX2* and *NEUROD2*; and *IL1RAPL1*, modulated by *SOX2*, *ZNF219*, *CTCF* and *TBR1* (Fig. 2c, g). Both *PTPRD* and *IL1RAPL1* are associated with SCZ and depletion of both genes have been shown to induce aberrant neurogenesis, maturation, and behaviors in mouse models^72–74^. This finding highlights strong connections among disease-related genes in TF-driven regulatory networks, indicating their potential roles as hub effector genes downstream of developmental TFs.

## Distinct fate outcomes of human RG upon TF perturbations

We further investigated the impact of TF perturbations on cell type composition using cluster free approaches to map cell fate changes with higher resolution along developmental trajectories (Fig. 3a). These approaches yielded consistent results with cluster-aware methods at the gene and individual sgRNA level and were robust to downsampling (Fig. 3a, Extended Data Fig. 3e-f, Extended Data Fig. 4).

**Fig. 3:**
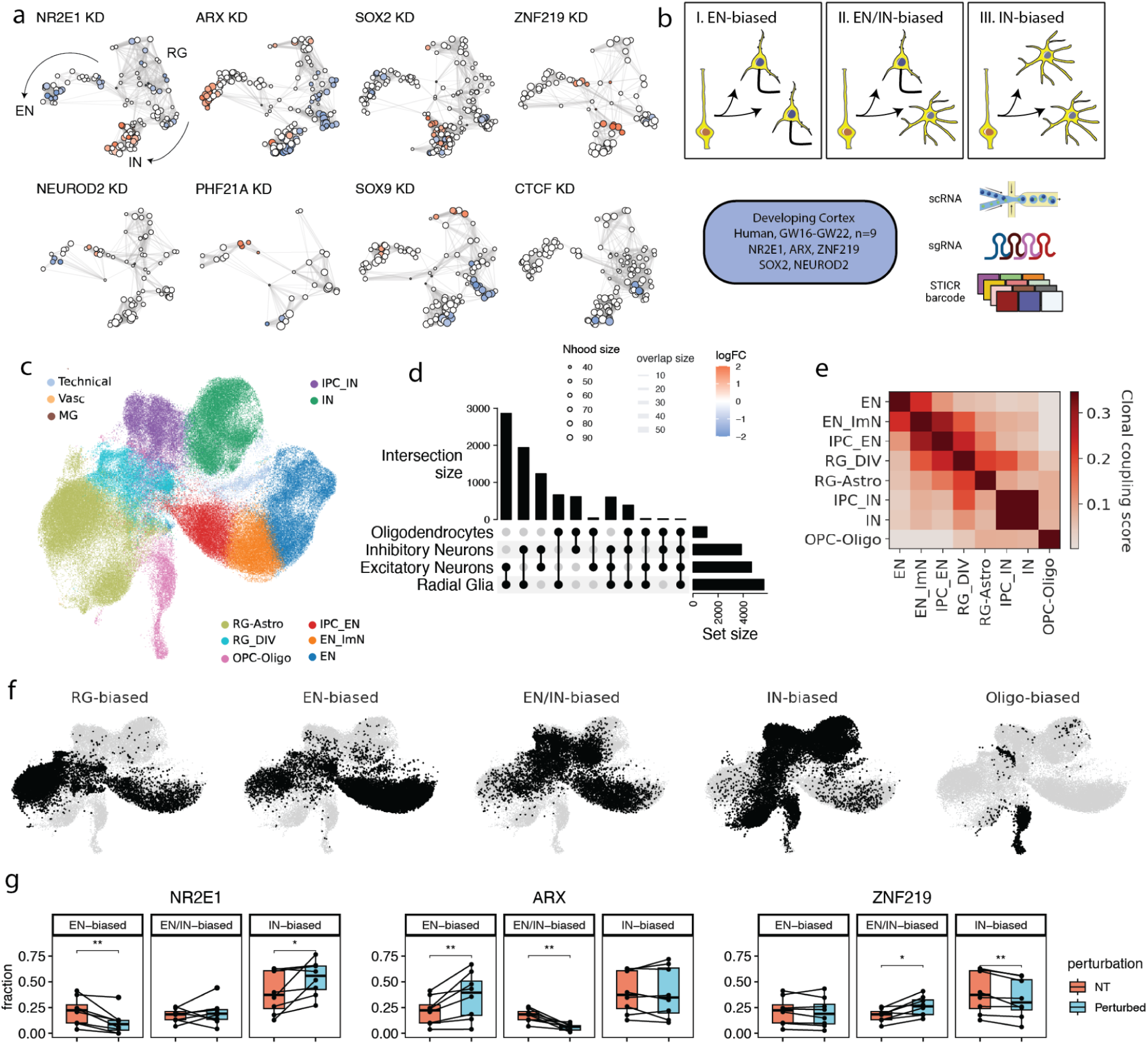
TF perturbations alter EN/IN output of human cortical RG. a. Neighborhood graphs based on cell positions in the Fig. 1d UMAP to show results of differential abundance testing using Milo^101^ under perturbation of 8 prioritized TFs at the gene level. NT cells were subset based on the number of perturbed cells to balance the cell number between conditions. Nodes are neighborhoods with sizes representing the number of cells in each neighborhood. The edges depict the number of cells shared between neighborhoods and the layout of nodes is determined by the position of the neighborhood index cell in the UMAP in Fig. 1d. The nodes are colored by their log_2_FC (FDR=0.1) when perturbed. b. Schematics for RG of distinct fate biases (top) and experimental design for combined TF perturbation and lineage tracing in cortical RG of 9 human individuals (bottom). c. Integrated UMAP of lineage-resolved 129,003 human cells collected on differentiation D7, colored by supervised cell type. A median of 15,469 cells per TF perturbation were recovered after filtering. d. Upset plot of cell class compositions in clones that have cells from more than one class (multi-class clones). e. Heatmap for lineage coupling score matrix in major cell types. f. UMAPs for cell fate distributions in clonal clusters identified using scLiTr^87^: RG-, EN-, EN/IN-, IN- and Oligo-biased clusters. g. Box plot of distributions of clonal cluster fractions in 8 human individuals in *NR2E1, ARX* and *ZNF219* KD in comparison to NT, grouped by clonal clusters. Significant changes under two-sided paired Wilcoxon tests were labeled. *, p<0.05; **, p<0.01; ***, p<0.001

Different perturbations elicited a range of composition consequences in differentiated cell types (Fig. 2b and Extended Data Fig. 3e-f). These phenotypes included examples consistent with mechanistic studies. For example, repression of SOX2 resulted in accumulation of RG^75^, while repression of the proneural factor, *NEUROD2* enriched for progenitors and immature ENs at the expense of mature ENs (Fig. 3a, Extended Data Fig. 3e). However, screening also revealed additional roles for previously characterized genes implicated in disease. For example, *NR2E1* knockout was previously shown to drive premature neural differentiation in mouse models^76,77^, but repression in human RG specifically enriched for postmitotic INs, in addition to depleting RG (Fig. 3a, Extended Data Fig. 3e). Similarly, *ARX* knockout was previously shown to affect IN differentiation and migration in mouse models^78–80^, but repression in human RG specifically enriched for the EN lineage at the expense of INs.

Our screen also uncovered disease-linked genes with composition phenotypes that to our knowledge have yet to be functionally examined during cortical neurogenesis, including *ZNF219* and *PHF21A*. Repression of *ZNF219* mirrored the effects of repressing *SOX9*, a known ZNF219 interaction partner in chondrocytes^81^, by increasing the proportion of ENs, but showed different effects from *SOX9* in dividing RG and INs (Fig. 3a, Extended Data Fig. 3e). Similarly, *PHF21A*, a member of the BRAF/HDAC complex implicated in synapse formation^82^, is a causal gene for ASD, along with *ARX* and *CTCF*^83,84^. Targeting each of these three genes led to an overall trend towards EN over IN differentiation (Fig. 3a, Extended Data Fig. 3e,f).

Flow cytometry analysis of the top TF candidates, *ARX, NR2E1* and *ZNF219*, with finer temporal resolution through flow cytometry validated the cellular phenotypes observed in our Perturb-seq (Extended Data Fig. 5, Supplementary Table 6), and further revealed early manifestations of compositional changes in KI67+ and EOMES+ progenitors on differentiation D4, confirming that these perturbations affect self-renewal and cell fate decisions at the stage of neurogenesis instead of neuronal maturation (Extended Data Fig. 5d).

## TF perturbations alter EN to IN output of individual RG

The influence of multiple TFs on the relative abundance of ENs and INs nominated RG lineage plasticity as a candidate developmental mechanism for observed composition differences. Composition differences could arise from perturbations of RG that preferentially affect EN biased clones, mixed clones containing both ENs and INs, or IN biased clones (Fig 3b). To distinguish between these possibilities, we combined Perturb-seq with lineage tracing using STICR, a GFP-expressing lineage tracing library with static barcodes that can be measured by scRNA-seq^1^. We constructed an mCherry-expressing sub-library targeting *NR2E1*, *ARX* and *ZNF219*, and also included *SOX2* and *NEUROD2* as controls, the former known to impact both lineages and the latter known to promote the EN lineage (Fig. 3b).

As RG fate plasticity changes with maturation, we extended the experiments using this targeted dual library to primary human RG culture isolated from nine human individuals from GW16 to GW22, spanning the peak of excitatory and inhibitory neurogenesis^12^ to early stages of gliogenesis. Cells co-expressing mCherry and GFP were isolated on D7 for scRNA-seq, and transcriptomes were reference-mapped to the developmental cell atlas described in Wang et al.^12^ and integrated with the initial Perturb-seq data for cell type annotation. This analysis yielded 129,003 cells with an assigned sgRNA and lineage barcode (Fig. 3c, Extended data Fig. 6a-j and Methods) and recapitulated the gene expression and cell type composition consequences observed in the initial screen (Extended data Fig. 6k,l).

We next investigated the landscape of clonal lineage relationships. We considered multicellular clones containing at least 3 cells with consistent sgRNA assignment, recovering 8,937 clones (Extended Data Fig. 7a and Methods). One human individual with low overall cellular coverage was dropped from this analysis. Remaining clones contained a mean of 5 cells in NT conditions (Extended Data Fig. 7b). The most abundant multi-class clones contained RG and ENs (35%) and RG and INs (20%), but, consistent with recent studies, we also observed around 15% clones containing both ENs and INs (Fig. 3d). Lineage coupling, defined as the normalized barcode covariance between cell types^85,86^, further supported the presence of mixed clones with a comparable linkage of dividing RG between both excitatory and inhibitory IPCs and immature neurons (Fig. 3e). Unsupervised clustering of clones based on cell type composition using scLiTr^87^ (Methods) revealed five categories of fate biases, comprising RG, EN, IN, oligodendrocytes, and EN/IN dual fate biased clonal clusters (Fig. 3f; Extended Data Fig. 7g). Interestingly, a small percentage of INs were observed in oligodendrocyte-biased clones and vice versa, suggesting their close lineage relationship^12,88^.

Intersecting perturbation and lineage tracing information revealed fate alterations by different TF perturbations. *NR2E1* KD significantly enriched for IN-biased clones at the expense of RG- and EN-biased clones, while *ARX* KD strongly enriched for EN-biased clones at the expense of EN/IN dual fate biased clones (Fig. 3g, Extended Data Fig. 7h). A significant decrease in the fraction of EN/IN dual fate biased clones was also observed upon *ZNF219* KD, suggesting that it shapes the balance of EN/IN output by increasing the diversity of lineage output from individual RG. In each case, the alterations in RG lineage biases were consistent with the observed cell type composition changes (Fig. 3a, Extended Data Fig. 6k), supporting lineage plasticity as the developmental mechanism underlying the effects of these TFs on EN and IN fates.

## TF perturbations alter RG lineage progression

RG undergo temporal fate restriction sequentially producing distinct subtypes of neurons as well as glial cell types. Along the developmental axis, we observed a stage-dependent shift in RG fate biases marked by a decrease of EN-biased clones and increase of IN-clones around GW18-19^12^ (Fig. 4a), supported by increased lineage coupling between dividing RG and IN after GW19 (Extended Data Fig. 7i). Interestingly, at GW19, *ARX* KD delayed the timing of IN-biased cluster expansion, while *NR2E1* KD promoted this transition (Fig. 4a), suggesting stage-dependent effects of TF perturbations on lineage decisions. Furthermore, there appeared to be a critical time window for altering EN/IN fate before GW21-22 (Extended Data Fig. 7h). Pseudotemporal ordering based on clonal types recapitulated the sequence of temporal cell fate choices beyond the EN/IN transition (Fig. 4b-e). In addition to increasing IN-biased clones at early stages, we found that *NR2E1* KD also increased Oligo-biased clones at later stages, nominating this TF as a regulator of developmental tempo (Fig. 4e-f, Extended data Fig. 7h). Conversely, *ARX* KD significantly shifted the clonal pseudotime distribution to more immature stages. Together, analyzing lineage progression across differentiation stages, revealed opposing effects of *NR2E1* and *ARX* on delaying and accelerating RG lineage progression, respectively (Fig. 4f).

**Fig. 4:**
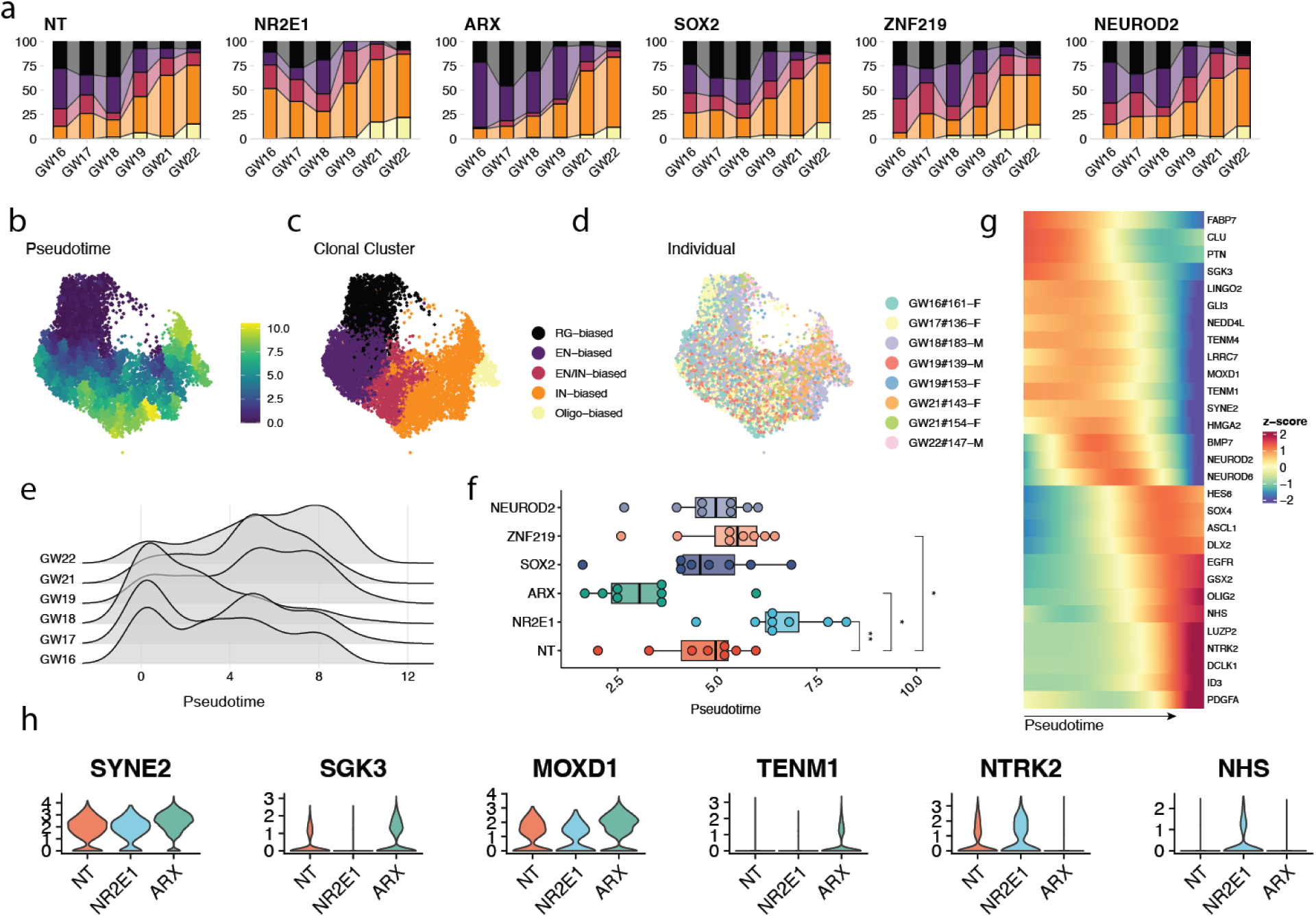
Heterochronic RG lineage progression under TF perturbations. a. Stacked barplot of distributions of clonal cluster fractions along human developmental stages in NT, *NR2E1, ARX, SOX2, ZNF219 and NEUROD2* KD, colored by clonal clusters. Legend is shown in Fig. 4c. b-d. UMAPs of human clones colored by (b) pseudotime, (c) fate biases and (d) individuals. e. Ridge plots of pseudotime distribution in clones grouped by stages. f. Boxplot to show median value of clonal pseudotime in each individual under TF perturbation, highlight opposing roles of *ARX* and *NR2E1* in regulating maturation of cell fate potential. Paired two-sided Wilcoxon test was performed to test for significance. g. Heatmap to show changes in gene expression in RG along the clonal pseudotime shown in Fig. 4b. h. Violin plots to show changes in gene expression in RG under *ARX* and *NR2E1* perturbations. *p<0.05; **, p<0.01; ***, p<0.001

To identify candidate effector genes involved in RG lineage progression, we compared gene expression in RG with different lineage histories along clonal pseudotime. Lineage tracing studies in rodents and gene expression data in humans have suggested that the fate transition of RG from EN to olfactory bulb (OB) IN neurogenesis involves a shift from GLI3 to EGFR signaling^12,88–90^. Similarly, RG in EN biased clones preferentially expressed *GLI3*, while those involved in IN production (EN/IN- and IN-biased clones) highly expressed *EGFR* and *PDGFRA* (Fig. 4g, Extended Data Fig. 7j). As the INs in these clones mainly expressed cortical-like markers including *SCGN* and *PAX6* (Fig. 1f, Extended Data Fig. 1f), this observation suggests reuse of a conserved signaling pathway transition for OB and local IN production by cortical RG. We further identified downstream genes under *NR2E1* and *ARX* KD with lineage dependent expression (Fig. 4h), nominating shared effector gene programs contributing to the RG lineage progression.

To investigate conservation of RG lineage plasticity and underlying mechanisms among primates, we extended our screening approach to rhesus macaque RG (Extended data Fig. 6d). The production of local INs by cortical RG has been suggested in rhesus macaque^91–93^, but not yet demonstrated by lineage tracing. We observed conservation in EN/IN lineage plasticity (Extended data Fig. 7e,f), including the production of cortical-like INs expressing *SCGN* and *PAX6*, and conserved changes in cell type compositions (Extended data Fig. 6k,m), gene expression (Extended data Fig. 6n,o, Supplementary Table 7) with a minority of divergent changes reflecting gene network evolution (Extended data Fig. 6p), and conserved effects of *ARX* and *NR2E1* on heterochrony of RG clonal output (Extended data Fig. 7k). Collectively, these results suggest conserved lineage plasticity of cortical RG in cortical-like IN neurogenesis across primates and conserved roles of *ARX* and *NR2E1* in modulating cortical developmental tempo.

## *ARX* safeguards postmitotic IN identity by repressing *LMO1*

Beyond lineage plasticity and progression, we also investigated the roles of TFs in subtype specification of cortical neurons. Both *ARX* and *SOX2* KD drove local composition changes among INs (Fig. 3a), motivating us to further examine their effects on subtype abundance. Iterative clustering of INs revealed populations expressing subtype markers including *ST18* (immature), *PAX6* (LGE-like), *CALB2* (CGE-like), as well as a small population expressing *PBX3* (OB-like) (Fig. 5a,b, Extended Data Fig. 8a,b). Interestingly, we observed two distinct subclusters exclusively defined by perturbations (“ectopic”): *ARX* KD created a cluster distinguished by *LMO1*, *a* previously identified target gene repressed by *ARX* in ventral telencephalon^71,78^ and *RIC3*, a chaperone for nicotinic acetylcholine receptors (Fig. 5b,c), while *SOX2* KD created a subcluster highly expressing *SPOCK1*, a neuronally produced and secreted proteoglycan (Extended Data Fig. 8c and Supplementary Table 5). Consistent results were observed across multiple sgRNAs sharing the same target, between individuals, batches and species (Extended Data Fig. 8d-f), highlighting conserved roles of *ARX* and *SOX2* in not only regulating neurogenesis but also specifying normal IN subtype identity, as well as the distinct ectopic transcriptional states formed by their loss.

**Fig. 5:**
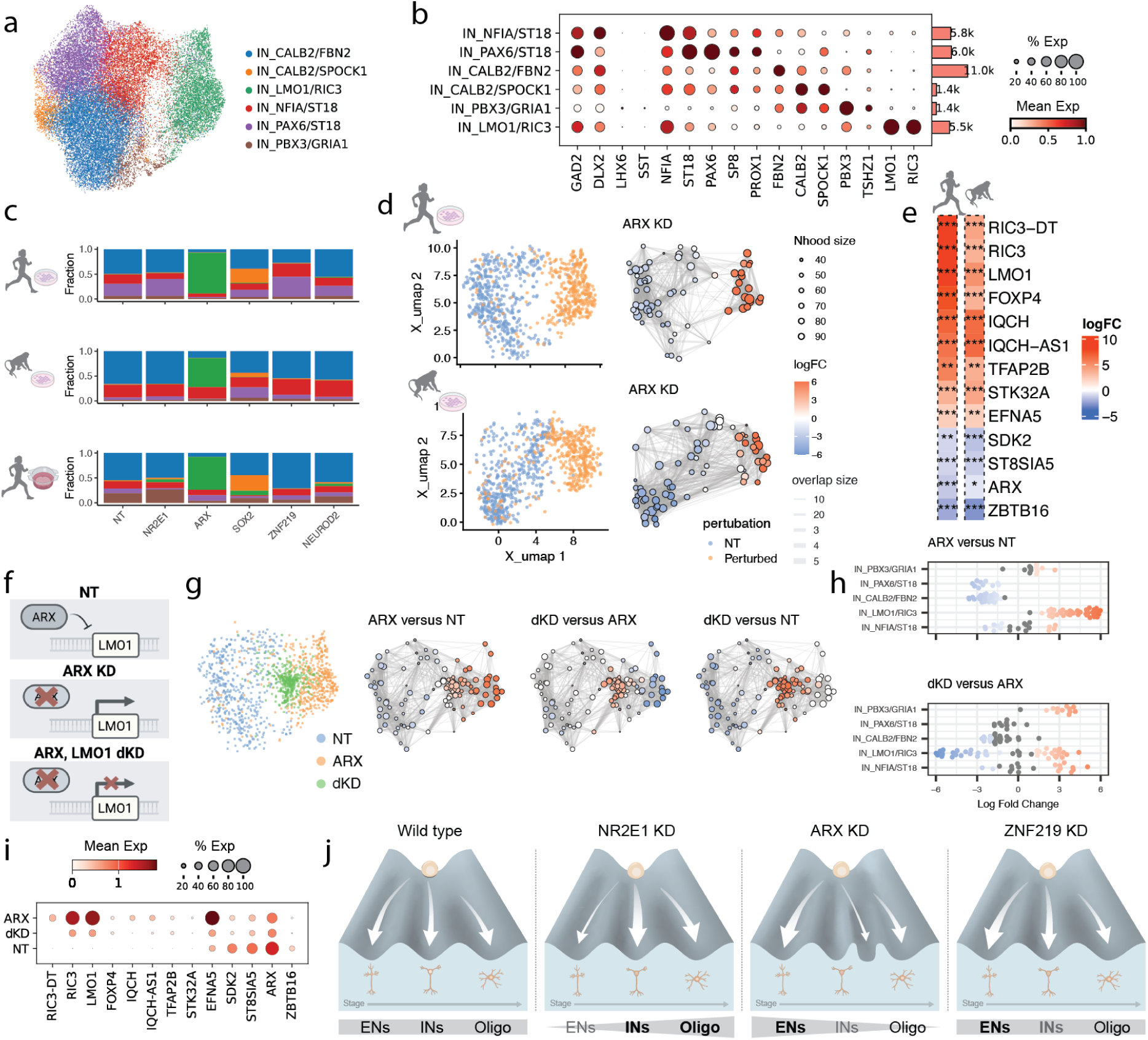
ARX safeguards IN identity by repressing LMO1. a. UMAP of integrated human and macaque IN clusters identified in the lineage resolved 2D screen. b. Dotplot showing marker gene expression in each IN cluster (left), with a barplot showing numbers of cells detected in the cluster (right). The size of each dot denotes the fractions of cells in the group where the gene is expressed and the color denotes mean gene expression in the group. c. Stacked barplot of distributions of IN subclusters in each TF perturbation in human (top) and macaque (middle) 2D and in human slice culture (bottom), colored by IN subtypes. d. UMAPs for differential abundance testing in IN subtypes using Milo in ARX KD at the gene level in human (top) and macaque (bottom). NT cells were subset based on the number of perturbed cells to balance the cell number between conditions. (left) UMAP of subset NT and perturbed cells from D7, colored by perturbation condition and labeled by IN subcluster. (right) Neighborhood graph of differential abundance testing colored by log_2_FC under FDR=0.05. e. Heatmaps of log_2_FC in top DEGs in IN_*LMO1/RIC3* cluster compared to other IN subtypes in human (left) and macaque (right). Dot/asterisks indicate significant changes identified from DEseq2. f. Schematics for gene expression regulation under NT, *ARX* KD and *ARX, LMO1* double KD (dKD). g. UMAPs for differential abundance testing in IN subtypes using Milo in NT, *ARX* KD and dKD. (left) UMAP of subset NT and perturbed cells colored by perturbation conditions. (right) Neighborhood graph of differential abundance testing colored by log_2_FC (FDR=0.05) under different contrasts: *ARX* KD versus NT; dKD versus *ARX* KD and dKD versus NT. h. Beeswarm plots showing differential abundance in neighbourhoods identified in Fig. 5g at the IN subtype level. Neighbourhoods that had significant changes between conditions were colored by their log_2_FC. i. Dotplot showing gene expression changes in IN_*LMO1/RIC3* marker under *ARX*, *LMO1* double KD. j. Graphical summary of findings: Human cortical RG gives rise to EN, IN and oligodendrocytes sequentially along developmental stages. NR2E1 KD promotes the RG lineage progression while ARX KD delays the transition and impairs normal IN subtype specification. ZNF219 KD promotes differentiation of both EN and IN with an overall preference to EN. •, p <0.1, *, p<0.05; **, p<0.01; ***, p<0.001

To further test the relevance of these findings to *in vivo* differentiation trajectories, we employed organotypic cortical slice culture which preserved the 3D organization of cortical tissue. To preferentially label progenitor populations we locally transduced germinal zone, where progenitors and immature cell types reside, with the targeted CRISPRi and lineage tracing libraries (Extended Data Fig. 9a-c). While the sensitivity of this model for detecting cell fate consequences is limited by heterogeneous starter cells and spontaneous differentiation in advance of gene repression^39^, the slice culture experiment recapitulated the transcriptomic changes observed in 2D culture (Extended Data Fig. 9d,e) and enabled studies of subtype specification. We observed immature cortically derived INs displaying low expression for migration markers *ERBB4* and *CXCR4* and lineage coupling to ENs (IN_local), along with mature CGE- and medial GE (MGE)-derived IN clusters that were likely labeled post-mitotically and not coupled to ENs (Extended Data Fig. 9f,g). Among INs, ARX repression again induced the *LMO1/RIC3* subtype, while SOX2 repression induced the *CALB1/SPOCK1* subtype, as observed in the 2D model (Fig. 5c, Extended Data Fig. 9h).

Differential gene expression analysis focusing on the *LMO1/RIC3* subtype revealed dysregulated activin and TGF-B signaling pathways driven by *ACVR1, SMAD2/3*, as well as impaired cell migration (*SMAD3, FOXP4, CDH20*) and synaptic membrane adhesion (*PTPRD, EFNA5*) (Supplementary Table 8). We observed consistent patterns of dysregulated genes in organotypic slice culture and conservation in rhesus macaque, consistent with previous reports of migration defects of INs in *ARX* mutant mice^56^ (Fig. 5e, Extended Data Fig. 8g-i). Despite dysregulation of genes in these pathways, we did not observe an elevated cellular stress response (Extended Data Fig. 8j).

To examine the developmental origin of these ectopic clusters, we analyzed lineage coupling and infection during differentiation. We observed similar lineage coupling across cell types in the RG lineage following *ARX* and *SOX2* perturbations in 2D culture (Extended Data Fig. 8k), suggesting that effects on subtype identity likely emerge post-mitotically rather than by altering initial progenitor fate specification. Further supporting a post-mitotic effect, the ectopic state emerged in cortical born INs in slice culture under conditions in which RG were not expanded and were permitted to differentiate immediately (Fig. 5c and Extended Data Fig. 9f,h). Additionally, we observed a similar ectopic cluster characterized by *LMO1* induction in CGE and MGE-derived INs that were infected post-mitotically (Extended Data Fig. 9j-l). Consistent with this observation, ARX expression is retained in post-mitotic INs (Extended Data Fig. 9m-o). Reference mapping to an *in vivo* atlas of cortical INs showed modest similarity between *LMO1/RIC3* subtypes and MGE derived *LHX6*+ INs, but missing canonical MGE marker expression suggests this subtype is unlikely to have a physiological counterpart (Fig. 5b and Extended Data 9p).

We next examined the molecular mechanisms underlying the emergence of the *LMO1/RIC3* cluster. As *LMO1* is a conserved downstream target of ARX across species whose expression is dysregulated in *ARX* mutant mice^71,94,95^ and acts as a transcriptional coactivator, we reasoned that *LMO1* may mediate the transcriptional programs driving the ectopic cell state. We performed double KD (dKD) with *ARX* and *LMO1* (Fig. 5f) by combining CRISPRi vectors expressing GFP and mCherry and integrated the resulting cell populations with previous data (Extended Data Fig. 10a-b). Reference mapping of IN subtypes revealed enrichment of an intermediate cell state between the *LMO1/RIC3* subtype and other subtypes, suggesting a partial rescue of ectopic cell state driven by *ARX* KD (Fig. 5g,h). Consistent with this model, *LMO1* dKD rescued the expression of many conserved marker genes identified in the *LMO1/RIC3* subtype, including *RIC3* and *EFNA5* (Fig. 5i and Extended Data Fig. 10c-e). These results indicate that *ARX* safeguards normal post-mitotic IN identity, in part by repressing *LMO1*, which is necessary for inducing additional genes characterizing the ectopic cluster. Together, our study introduces a primary cell model of cortical neurogenesis with improved fidelity to normal development and reveals a broadly conserved landscape of responses to TF perturbations in gene regulatory networks, cell fate decisions, and neuronal subtype specification (Fig. 5j).

## Discussion

Systematic profiling of single cell gene expression and chromatin accessibility during human cortical development has supported construction of human cell atlases, inference of developmental trajectories, and prioritization of gene regulatory networks^9,10,96,97^. However, functionally examining candidate regulators requires scalable approaches for perturbing gene expression in physiological models of human cortical neurogenesis. We established a primary culture system with improved fidelity and decreased cellular stress response, and applied single cell Perturb-seq in primary human and macaque RG undergoing differentiation to examine the role of 44 TFs in cortical development at the level of gene expression, gene network interactions, cell fate biases, and subtype specification. Convergence of effector genes downstream of multiple TFs and their enrichment in neuropsychiatric and neurodevelopmental processes and disorders highlighted the connectedness of TF networks during neurogenesis. Different perturbations elicited distinct RG fate outcomes. By coupling TF perturbation and barcoded lineage tracing, we showed the lineage plasticity of individual human and macaque RG. We identified *ZNF219* as a novel regulator in human cortical neurogenesis, and *NR2E1* and *ARX* as known regulators with new functions in regulating inhibitory neurogenesis and guiding temporal lineage progression of individual cortical RG, with postmitotic roles for *ARX* in IN subtype specification, partially through repressing *LMO1*.

RG undergo waves of neurogenesis and gliogenesis, sequentially generating distinct subtypes of neurons, oligodendrocytes and astrocytes. We derived a culture system that sensitively captured the fate transition of EN to IN production and showed the lineage plasticity of individual RG in response to TF perturbations. *ARX* repression extended the window of EN production, while *NR2E1* repression accelerated the transition to IN and oligodendrocyte production. Mapping clonal clusters along the developmental axis revealed variation in effect sizes of the fate switch potential in RG at different maturation stages, suggesting a restricted time window (GW18-19) for lineage plasticity, corresponding to the timing of endogenous EN/IN transition^12^. Combining perturbations with multimodal profiling including chromatin accessibility of RG at different stages could reveal the underlying *cis*-regulatory changes that account for different cell fate and developmental tempo responses downstream of TF depletion, while extending to genome scales could reveal additional regulators of protracted maturation of human RG.

The dorsal origin of a subset of human cortical inhibitory neurons has been supported through lineage tracing using both lentiviral-based static barcodes and somatic mosaicism^1,12,25,26^, but underlying mechanisms regulating this process remained elusive. Previous studies highlighted RG lineage plasticity as a potentially human-specific process in comparison to mouse^1,24^, but our lineage tracing experiments in macaque revealed conservation of lineage plasticity and responses to TF perturbations among primates. Further studies of IN migration and molecular identity will help illuminate possible quantitative differences between species and to distinguish between cortical and OB-bound INs generated by cortical RG.

Beyond cell fate commitment and maturation, our screening platform robustly detected changes in IN subtype specification, exemplified by conserved phenotypes observed in *ARX* and *SOX2* KD across primates. Pathogenic ARX mutations associated with neurologic disorders show altered DNA binding preferences and loss of *LMO1* repression^94,95^, consistent with the *LMO1* as a defining marker of the ectopic cell type. Dual repression of ARX and its downstream target *LMO1* partially reverted transcriptional programs driving the *ARX*-mediated ectopic IN state, highlighting *LMO1* repression as crucial for normal IN identity. Increased *LMO1* expression has been associated with tumor invasion and metastasis^98,99^, highlighting its potential role in ectopic IN migration. While *Ebf3,* another *Arx* target, has previously been implicated in Arx-mediated migration deficits^100^, future studies combining time lapse imaging and human cellular models can examine the role of *LMO1* as a master regulator mediating aberrant cell behaviors. The observation that the TFs with the strongest transcriptional and cell composition consequences have also been implicated in neurodevelopmental disorders highlights the correspondence of cellular phenotypes in a simplified culture system to organismal consequences. Collectively, our study provides a framework for functional dissection of gene regulatory networks in human cortical neurogenesis.

## Acknowledgments

The authors thank M. Paredes, D. Shin, M. Steyert, L. El Hayek, N. Sanchez-Luege and other Pollen lab members for valuable comments, N. Schaefer for supporting sgRNA assignments, J. Srivastava and V. Saware for their assistance performing cell sorting at the Gladstone Flow Cytometry Core, supported by NIH S10 RR028962, L. Wang for access to the multiome dataset in the developing human brain, S. Wang for transferring samples, and A. Tarantal for providing samples. Sequencing was performed at the UCSF CAT, supported by UCSF PBBR, RRP IMIA, and NIH 1S10OD028511-01 grants. Figures were made, in part, with BioRender. This work was supported by the following funding sources: CIRM fellowship (J.W.D.), National Institutes of Health R01MH134981-01, DP2MH122400-01 (A.A.P.), UM1MH130991 (A.A.P., T.J.N.), R01NS123263 T.J.N., Schmidt Futures Foundation (A.A.P., T.J.N.), William K. Bowes Jr. Foundation (A.A.P and T.J.N.). Shurl and Kay Curci Foundation (A.A.P., T.J.N.), the California Institute for Regenerative Medicine (CIRM) DISC0-14429 (T.J.N.) DISC4-16285 (A.A.P), as well as gifts from Esther A. & Joseph Klingenstein Fund. T.J.N. is a New York Stem Cell Foundation Robertson Neuroscience Investigator. A.A.P. is a New York Stem Cell Foundation Robertson Investigator.

## Contributions

J.W.D. and A.A.P. conceived the project and experimental design. A.A.P. supervised the study and secured the funding. J.W.D. performed all the experiments and analyzed the data with the help of C.N.K. under the supervision of T.J.N. and A.A.P.. C.N.K. performed STICR barcode assignment. M.S.O. helped on the validation with flow cytometry. J.L.W. and B.J.P. performed initial testing of the primary culture system. Y.A. prepared and tested the CRISPRi vector. J.W.D. and A.A.P. prepared the manuscript with input from all authors.

## Ethics declarations

All studies were approved by UCSF GESCR (Gamete, Embryo, and Stem Cell Research) Committee.

## Competing interests

The authors declare no competing interests.

## Extended Data Figures

**Extended Data Fig. 1:**
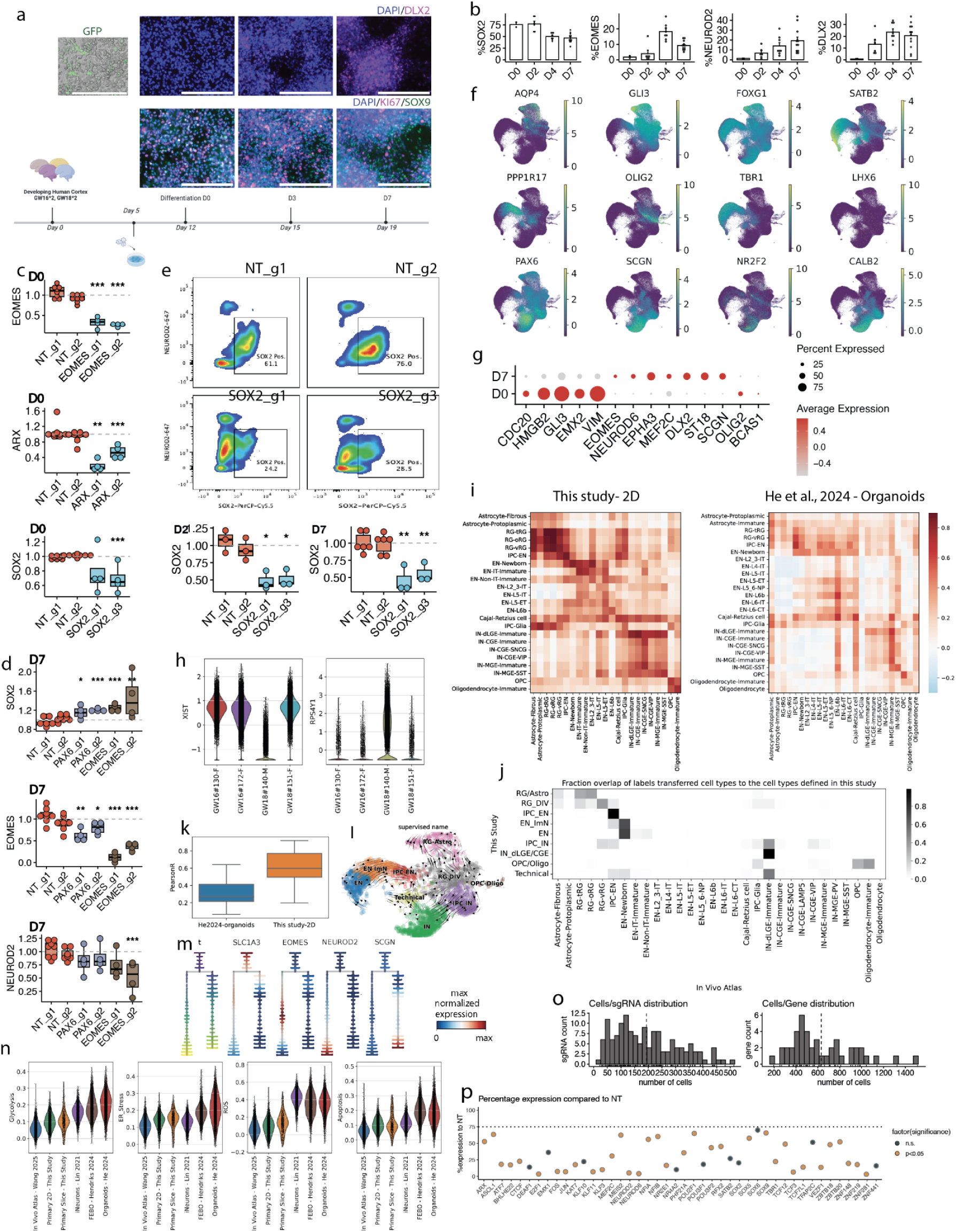
Supplemental analysis of the primary model, CRISPRi, and Perturb-seq data. a. *In vitro* culture of human cortical RG before (D0) and after induced differentiation (D3, D7), with immunocytochemical labeling of SOX9, MKI67 and DLX2. The top plot shows live cell imaging with GFP expression before differentiation. b. Barplot of SOX2, KI67, EOMES and DLX2 positive populations across the differentiation time course (D0, D2, D4, D7) in cells expressing all-in-one CRISPRi vectors with NT sgRNAs. Quantification was done through flow cytometry. Dots represent independent biological replicates and sgRNAs. D0, n=2; D2, n=3 x 2 sgRNAs; D4, n=3 x 2 sgRNAs; D7, n= 6 x 2 sgRNAs. c. Barplot of fold changes in fractions of EOMES, ARX and SOX2 positive populations compared to NT in each sgRNA to confirm efficient KD in *EOMES*, *ARX* and *SOX2* KD before differentiation (D0) obtained from flow cytometry from n=4 individuals. Dots represent biological replicates from independent individuals and sgRNAs. Paired two-sided Wilcoxon test was performed on the mean of fold changes compared to NT in each sgRNA, using individual replicates and NT as reference after collapsing 2 independent NT sgRNAs. d. Barplot of fold changes in fractions of SOX2, EOMES and NEUROD2 expressing populations obtained from flow cytometry in *PAX6* and *EOMES* KD at the sgRNA level in comparison to NT on D7 from n=4 individuals x 2 sgRNAs per perturbation. Dots represent biological replicates from independent individuals and sgRNAs. Increase in SOX2 and decrease in EOMES and NEUROD2 populations were detected in both perturbations under paired two-sided Wilcoxon tests. e. Flow cytometry data with quantified bar plot (bottom) of fraction of SOX2 positive populations compared to NT in each sgRNA to confirm efficient KD of SOX2 at the beginning (D2) and the end of differentiation (D7). Robust KD of SOX2 populations were detected throughout the course of differentiation under paired two-sided Wilcoxon tests. f. UMAP of cells collected at differentiation D0 and D7, colored by the expression of cell type markers. g. Dotplot of marker gene expression on D0 and D7. h. Violin plot of *XIST* and *RPS4Y1* expression in each individual, highlighting inferred sex genotype. i. Pearson correlation coefficients of top 25 cell marker gene expression defined in Wang et al^12^ *in vivo* atlas to data obtained in this study (left) and in He et al^53^ organoid atlas (right). Cell type annotation was done using reference mapping of both datasets to *in vivo* atlas in Wang et al^12^. j. Heatmap for fraction overlap of labels transferred from reference cell types^12^ to the cell types defined in this study. k. Boxplot to show distribution of Pearson correlation coefficients to predicted cell types (diagonal values from Extended Data Fig. 1h). l. UMAP of RNA velocity on NT cells collected on D7, colored by supervised cell types. m. Tree plot showing cells along pseudotime branches, colored by pseudotime and marker gene expression: *SLC1A3, EOMES, NEUROD2* and *SCGN*. n. Violin plots of gene expression scores of 5 types of cell stress (glycolysis, ER stress, ROS, apoptosis and senescence) in 6 different datasets including vivo human atlas, 3D organotypic slice culture, 2D primary RG culture, iNeurons^102^, primary brain organoids FeBO^52^ and iPSC-derived organoids^53^ obtained from published studies. Organotypic slice culture data presented at Extended Data Fig. 9 was used here for system benchmarking. o. Histogram for distribution of singly infected cell counts for each gene-targeting sgRNA (top) and TF (bottom) on D7. p. Dotplot for percentage target gene expression compared to NT cells for 44 TFs included in the study in the gene level after collapsing all active sgRNAs targeting the same gene. log_2_FC were calculated in each cell class and the lowest value for each gene was used for visualization and filtering. Dots are colored by p value and bordered based on log_2_FC of the target gene. ER, endoplasmic reticulum; ROS, reactive oxygen species Scale bars: 200 μm

**Extended Data Fig. 2:**
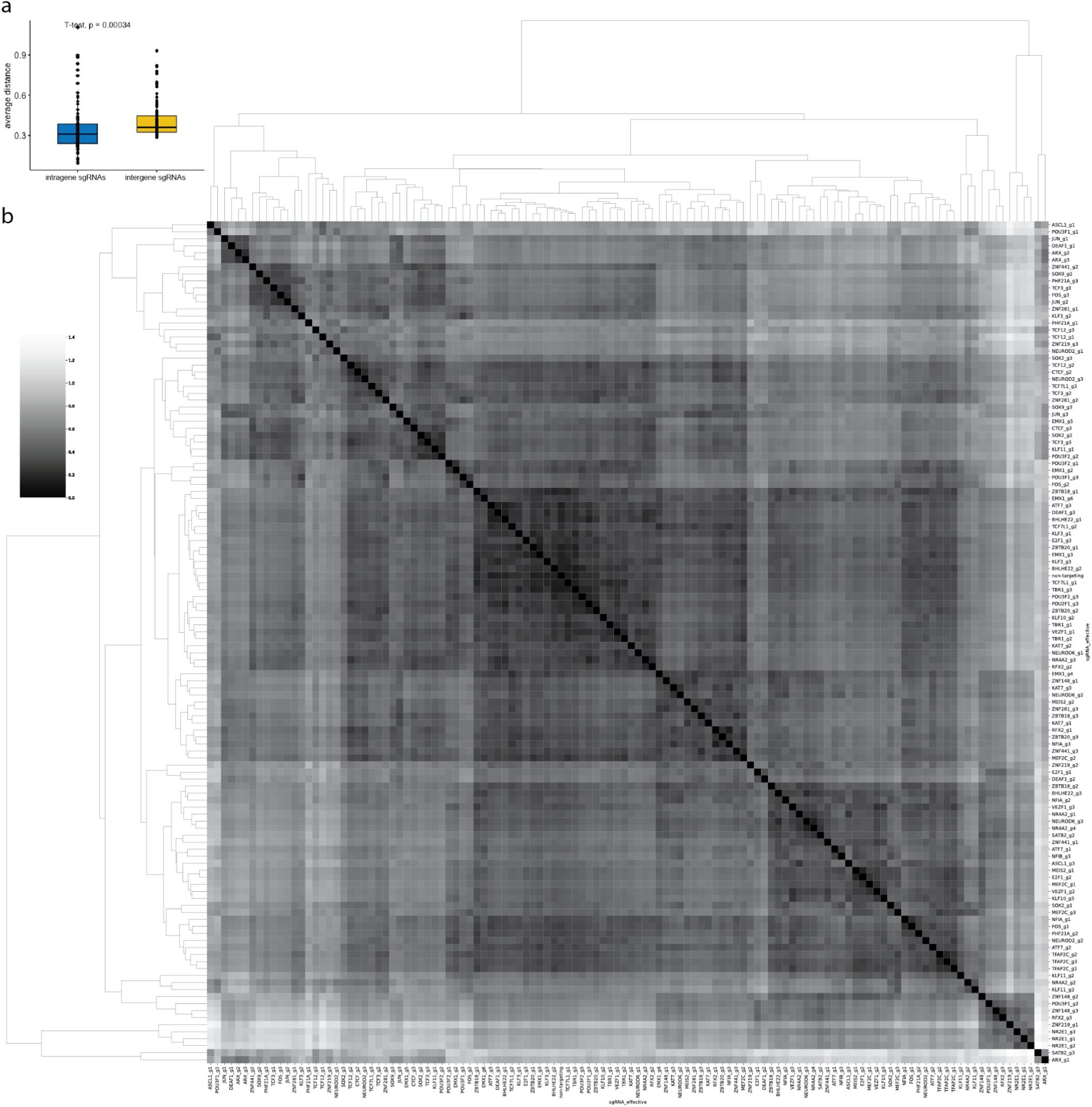
Euclidean distance between each sgRNA on differentiation D7. a. Boxplot for mean Euclidean transcriptional distance between each sgRNA with other sgRNAs targeting the same gene (intragene sgRNAs, blue) and different genes (intergene sgRNAs, yellow). Paired two-sided T-test was performed using each sgRNA as a replicate. b. Heatmap for Euclidean distance between each sgRNA.

**Extended Data Fig. 3:**
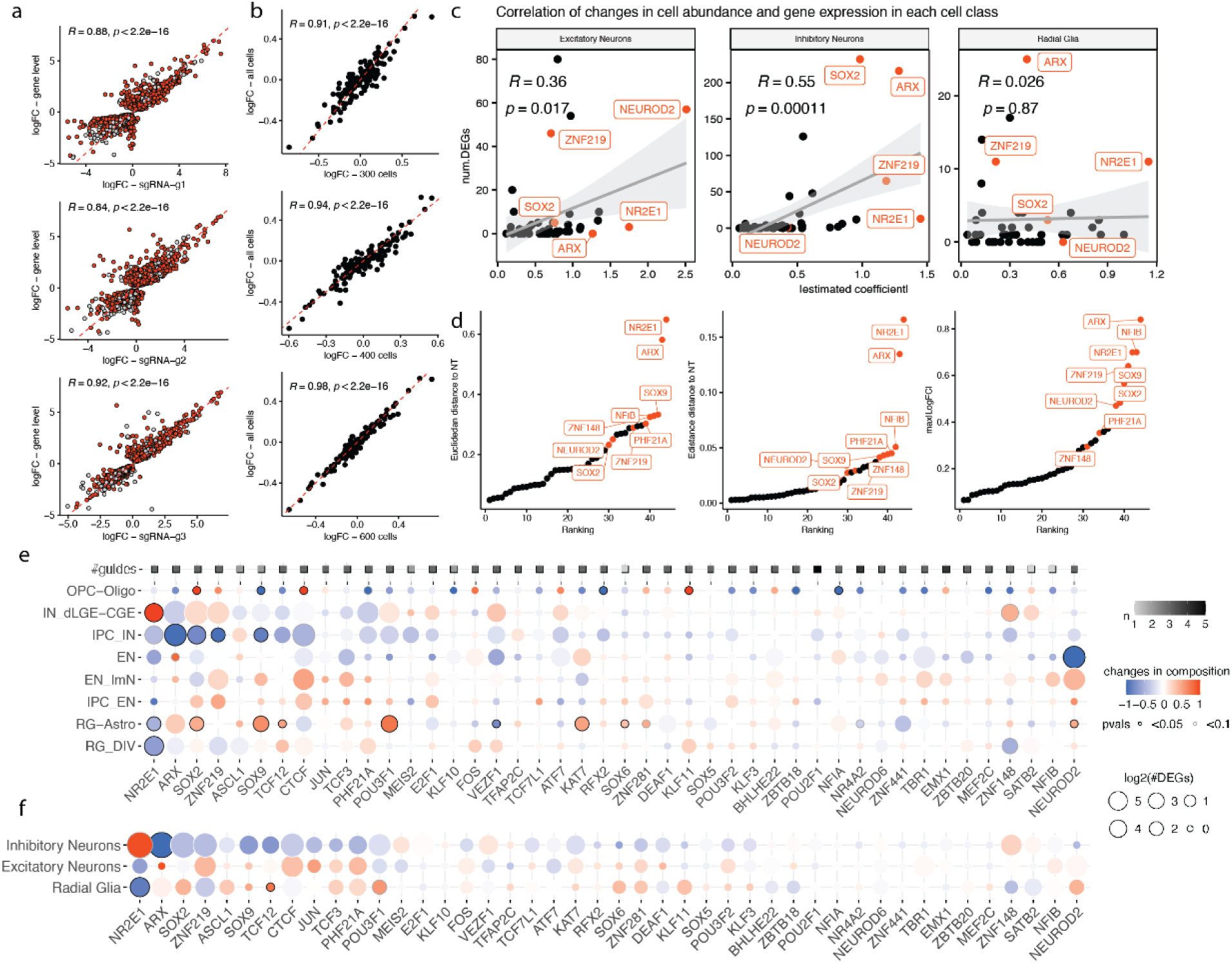
Supplemental analysis for responses to TF perturbation and target prioritization. a. Scatter plots of logFC of DEGs identified in each class in the sgRNA level (x axis) and gene level (y axis). Significant DEGs in the gene level were used for this comparison and red dots denote DEGs that were detected as significant in at least 2 conditions. Red dashed line represents Y=X. b. Scatter plots of logFC of cell abundance in each class in downsampled populations (x axis) and in all cells collected under each perturbation (y axis). Each condition was randomly downsampled to 300, 400 and 600 cells for this testing. Red dashed line represents Y=X. c. Scatterplot for summed absolute values of estimated coefficient in cell abundance (X axis) and number of DEGs (Y axis) in each cell type within 3 cell classes (Excitatory Neurons: IPC_EN, EN_ImN, EN; Inhibitory Neurons: IPC_IN, IN; Radial Glia: RG_DIV, RG-Astro) for each TF perturbation. *NR2E1, ARX, ZNF219, SOX2, NEUROD2* are labeled. Pearson correlation coefficients and p values were calculated in each cell class and labeled with linear regression lines with 95% confidence intervals. d. Scatterplot for (left) Euclidean distance to NT, (middle) energy distance^55^ to NT, (right) maximum absolute log_2_FC of cell abundance among 8 cell types listed in panel e calculated with scCoda^103^ at the gene level. e. Dot plot for global changes in cell abundance (color) and gene expression (size) with cell type resolution (rows) for 8 cell types under each TF perturbation (columns) on D7, with top row showing number of sgRNAs per TF used in this study after quality filtering. Fill color indicates estimated values of coefficients for differential abundance, calculated using DCATS^104^, with border intensity denoting p value from likelihood ratio tests; size indicates the number of DEGs, calculated using DEseq2 and filtered with adjusted p value < 0.05. f. Dot plot for global changes in cell type composition (color) and gene expression (size) in 3 broad cell classes under each TF perturbation. Note that the overall trend toward depletion of IN_IPCs across perturbations drives broader depletion of the IN lineage at the level of class, even in cases like *SOX2*, *ZNF219* which showed postmitotic IN enrichments.

**Extended Data Fig. 4:**
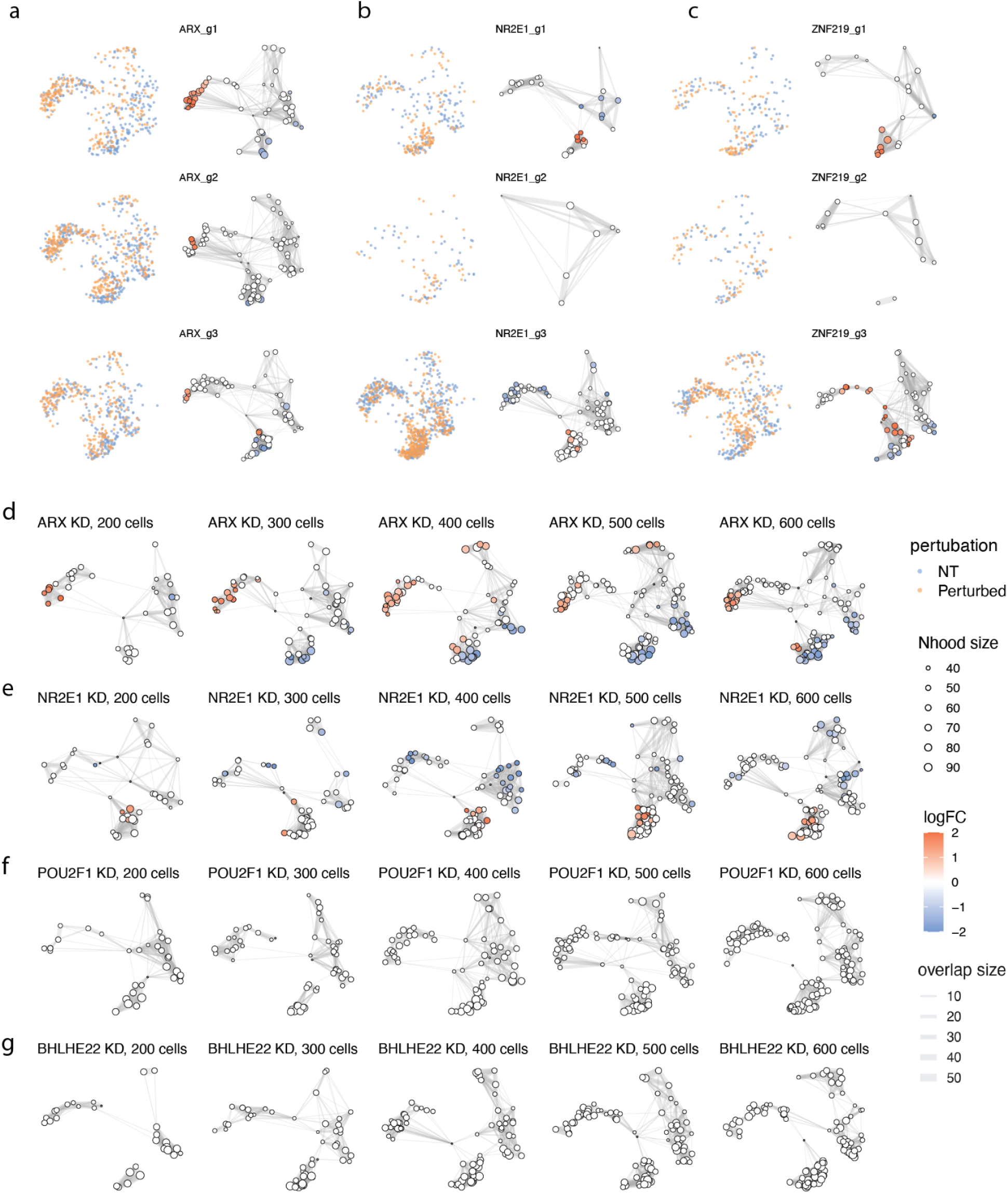
Supplementary analysis for cluster-free differential abundance testing using Milo. a-c. UMAPs of results for differential abundance testing using Milo^101^ at the sgRNA level for *ARX*(a)*, NR2E1*(b) and *ZNF219*(c) KD. (left) UMAP of subset NT and perturbed cells from differentiation D7, colored by perturbation condition. (right) Nodes are neighborhoods with sizes representing the number of cells in each neighborhood. The edges depict the number of cells shared between neighborhoods and the layout of nodes is determined by the position of the neighborhood index cell in the UMAP. The nodes are colored by their log_2_FC when perturbed. d-g. UMAPs of results for *ARX*(d), *NR2E1*(e), *POU2F1*(h), and *BHLHE22*(g) KD at the gene level after downsampling to 200-600 cells.

**Extended Data Fig. 5:**
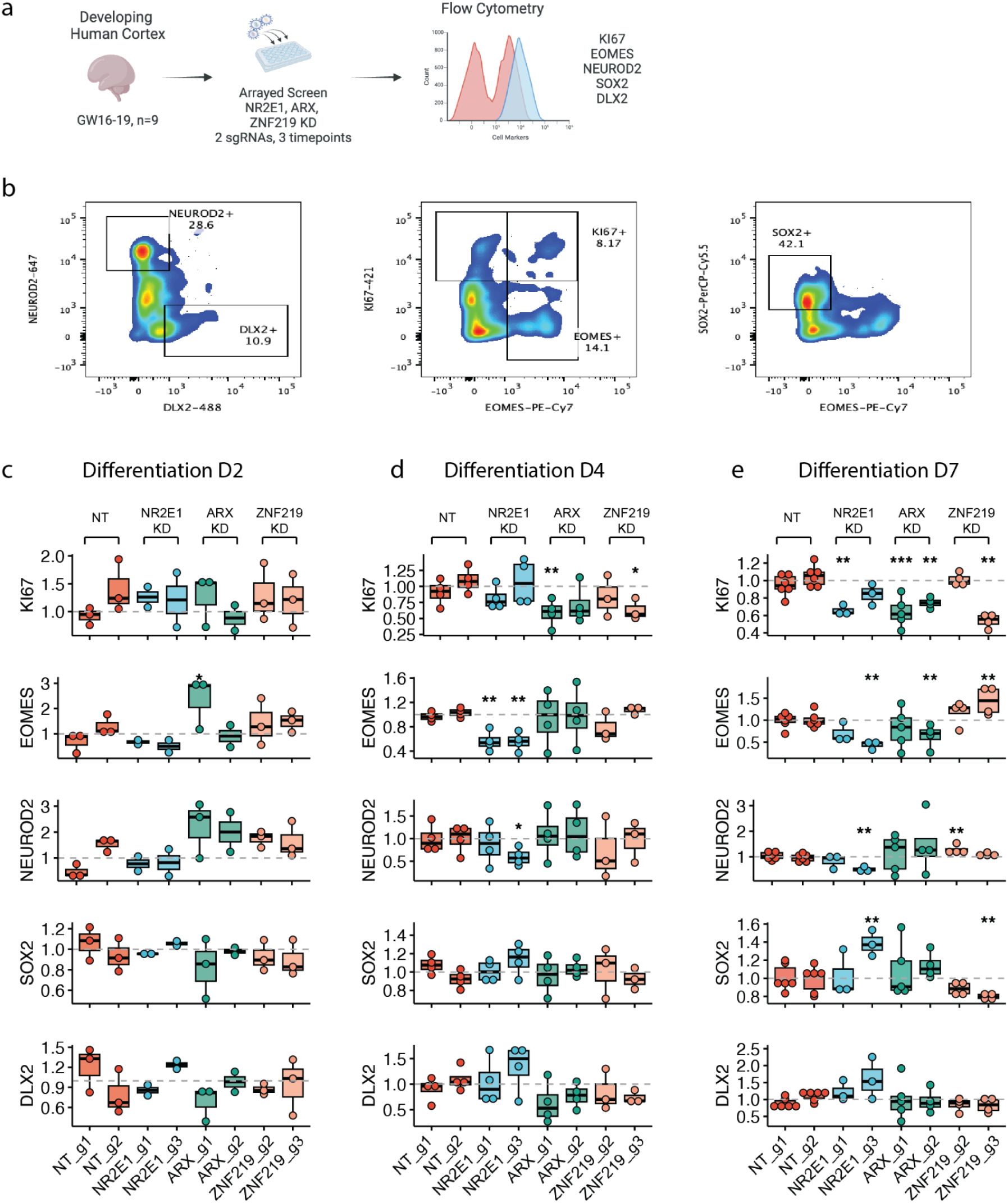
Cell type composition changes with temporal resolution during differentiation using flow cytometry. a. Experimental design of flow cytometry validation of prioritized targets from the Perturb-seq. The experimental workflow described in Fig. 1a was repeated on specimens of developing human cortical RG from 9 individuals spanning GW16-19 and fractions of KI67, EOMES, NEUROD2, SOX2 and DLX2 positive populations were quantified in mCherry-positive NT, *NR2E1*, *ARX* and *ZNF219* KD cells through flow cytometry. b. Example images of gating KI67, EOMES, NEUROD2, SOX2 and DLX2 positive populations on NT cells on differentiation D7. c-e. Barplot of fold changes in fractions of KI67, EOMES, NEUROD2, SOX2 and DLX2 expressing populations obtained from flow cytometry in *NR2E1, ARX* and *ZNF219* KD at the sgRNA level in comparison to NT. 2 sgRNAs were tested for each target on differentiation D2 (c), D4 (d) and D7 (e). Barplots are colored by the target gene and dots represent biological replicates from independent individuals. Paired two-sided Wilcoxon tests were performed on the mean of fold changes compared to NT in each sgRNA, using individual replicates and NT as reference after collapsing 2 independent NT sgRNAs. *, p<0.05; **, p<0.01; ***, p<0.001

**Extended Data Fig. 6:**
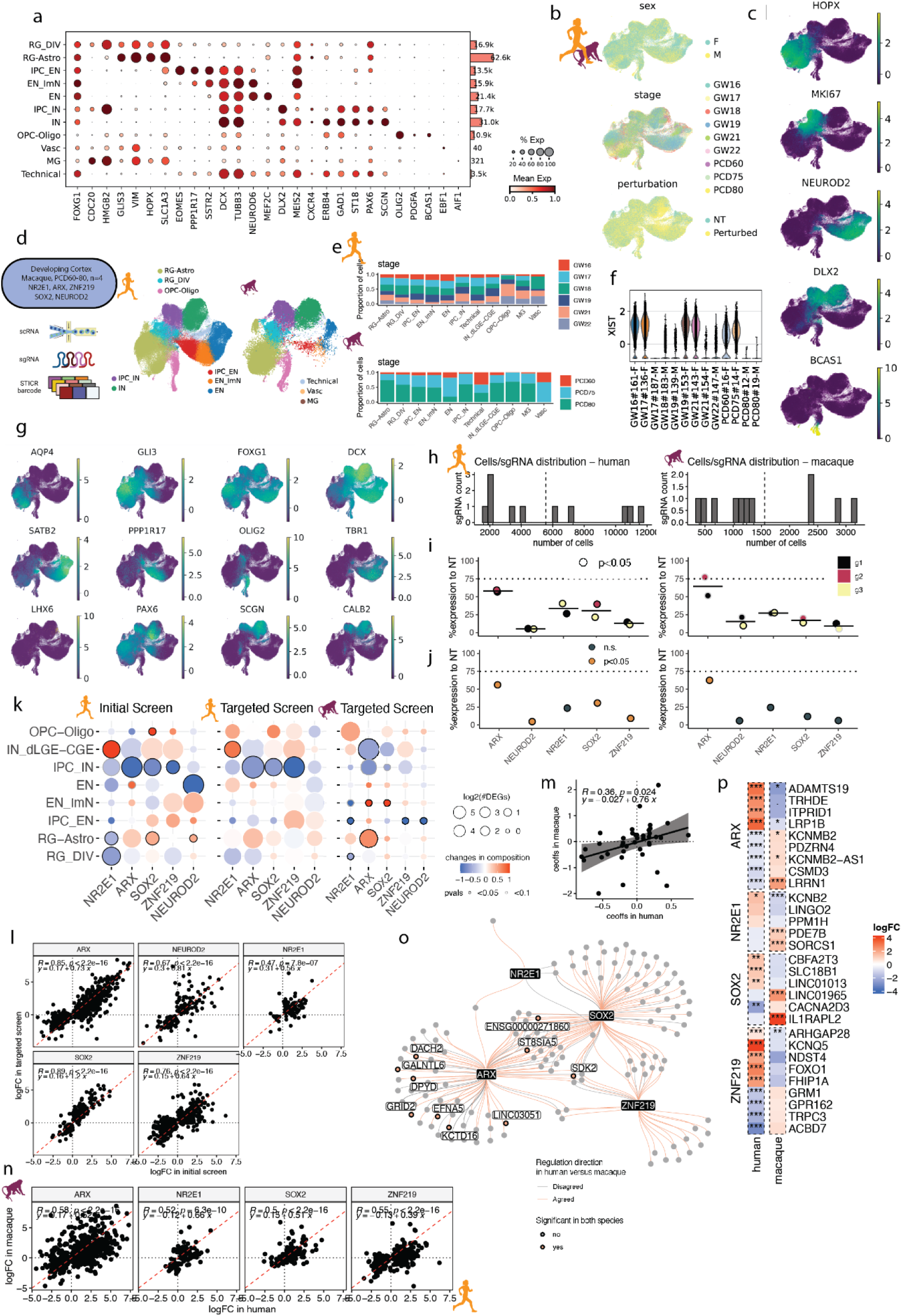
Lineage-resolved targeted Perturb-seq in humans and macaques. a. Dotplot of the expression of cell type markers for the assigned clusters (left), with a barplot showing numbers of cells detected in the cluster (right) in the lineage-resolved human and macaque dataset. The size of each dot denotes the fractions of cells in the group where the gene is expressed and the color denotes mean gene expression in the group. b. UMAP of integrated cells in both species, colored by sex, developmental stages and perturbation. c. UMAP of lineage-resolved 129,003 human (right) and 24,381 macaque (left) cells, colored by supervised cell types. A median of 15,469 and 3,013 cells per perturbation were recovered after filtering in human and macaque, respectively. d. UMAPs colored by the expression of lineage markers. e. Stacked barplot for distributions of developmental stages across cell types in human (top) and macaque (bottom). f. Violin plot of *XIST* expression in each individual, supporting inferred sex genotype. g. UMAPs colored by the expression of cell type markers. h. Histogram of the distribution of number of cells per sgRNA in the targeted library in human (left) and macaque (right). i. Dotplot for percentage target gene expression in each sgRNA compared to NT cells in human (left) and macaque (right) grouped by TF target, colored by sgRNA bordered based on significance. log_2_FC were calculated within each cell type and the lowest value for each sgRNA was used for visualization. j. Dotplot for percentage target gene expression in the gene level compared to NT cells in human (left) and macaque (right) grouped by TF target. Color indicates significance calculated by DEseq2 using individuals as biological replicates. k. Dot plot for global changes in cell abundance (color) and gene expression (size) with cell type resolution (row) under each TF perturbation (column) on differentiation D7 in the initial human screen (left), lineage-resolved targeted human (middle) and macaque (right) screen. Fill color indicates estimated values of coefficients for differential abundance, with border intensity denoting p-value from likelihood ratio tests; size indicates the number of DEGs and filtered with adjusted p value < 0.05. l. Scatterplot of log_2_FC for DEGs in the initial (X axis) and targeted (Y axis) human screens identified in *ARX, NEUROD2, NR2E1, SOX2* and *ZNF219* KD. Pearson correlation coefficients and p values were calculated in each TF perturbation and labeled together with regression line equations. Red dashed line represents Y=X. m. Scatter plot of estimated coefficients from DCATS in human (x axis) and macaque (y axis). Pearson correlation coefficients and p values were calculated in each TF perturbation and labeled together with regression line equations. n. Scatterplots of log_2_FC in human and macaque for DEGs identified in *ARX, NR2E1, SOX2* and *ZNF219* KD. Border intensity denotes disease association and DEGs that were significant in both species were colored based on regulation directions. Pearson correlation coefficients and p values were calculated in each TF perturbation and labeled together with regression line equations. Red dashed line represents Y=X. o. Regulatory network plot shows convergent DEGs (defined in Fig. 2) connected to *NR2E1, ARX, SOX2, ZNF219* in humans compared to macaques. Color of the edges indicates the conservation of regulation (pink suggests that DEG log_2_FC has consistent direction in 2 species). p. Heatmaps of log_2_FC in top divergent DEGs in response to *ARX, NR2E1, SOX2* and *ZNF219* KD in human and macaque. Dot/asterisks indicate significant changes identified from DEseq2. PCD, post conceptional day

**Extended Data Fig. 7:**
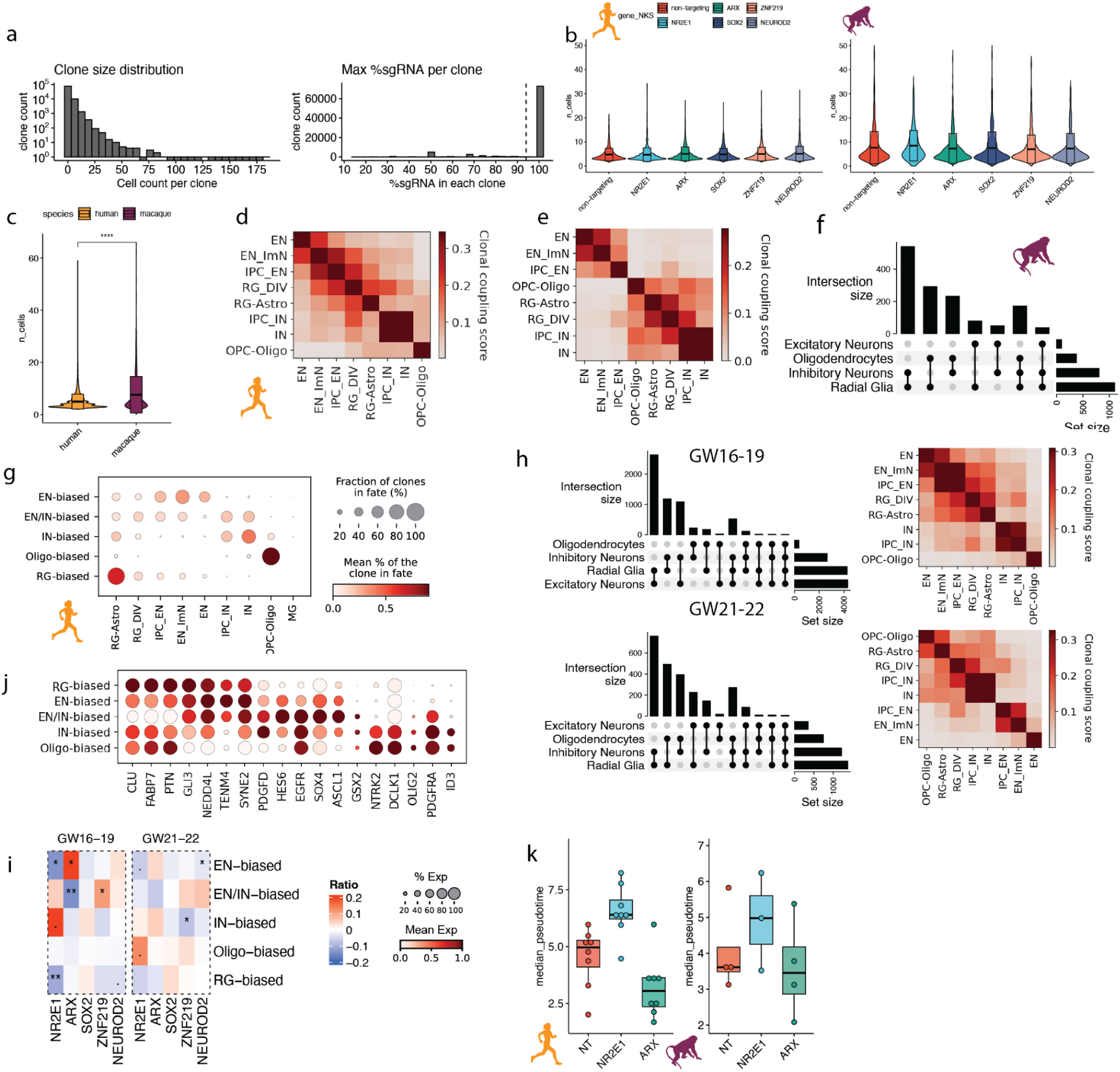
Supplemental analysis for RG lineage plasticity and progression. a. Histograms for distributions of (top) clone size and (bottom) maximum percentage of detected sgRNA in each clone in cells with assigned sgRNA before filtering. Clones with fewer than 3 cells or conflicted sgRNA assignments (maximum sgRNA percentage < 100%) were removed from downstream analyses. b. Violin plot for clone size distribution in human (left) and macaque (right), grouped and colored by TF target. c. Violin plot for clone size distribution in both species. Two-tailed T test was performed on the mean of clone sizes between species. d. Heatmap for lineage coupling score matrix in major cell types identified in human. e. Heatmap for lineage coupling score matrix in major cell types identified in macaque. f. Upset plot of cell class compositions in multi-class clones in macaque. g. Dotplot for distributions of supervised cell types in each clonal cluster identified in human. The size of each dot denotes the fractions of clones in each clonal cluster where the cell type is detected and the color denotes the mean percentage of each cell type in clones. h. Upset plots of cell class compositions in multi-class clones (left) and heatmaps for lineage coupling score matrix in major cell types (right) in human in early (top, GW16-19) and late (bottom, GW21-22) midgestation. i. Heatmap of changes in clonal cluster fractions under different perturbations in human in early (left) and late (right) midgestation. Color represents changes in fraction within each clonal cluster compared to NT cells. Dot/asterisks indicate significant changes identified from 2-sided paired T-test. j. Dotplots of marker gene expression in radial glia class (RG-Astro & RG_DIV) grouped by clonal clusters in human. The size of each dot denotes the fractions of cells in the group where the gene is expressed and the color denotes mean gene expression in the group. k. Boxplot showing median pseudotime of all clones in NT, *NR2E1* and *ARX* perturbation in human (left) and macaque (right). Dots represent biological replicates from independent individuals. Ctx, cortex; PFC, prefrontal cortex •, p<0.1; *, p<0.05; **, p<0.01; ***, p<0.001

**Extended Data Fig. 8:**
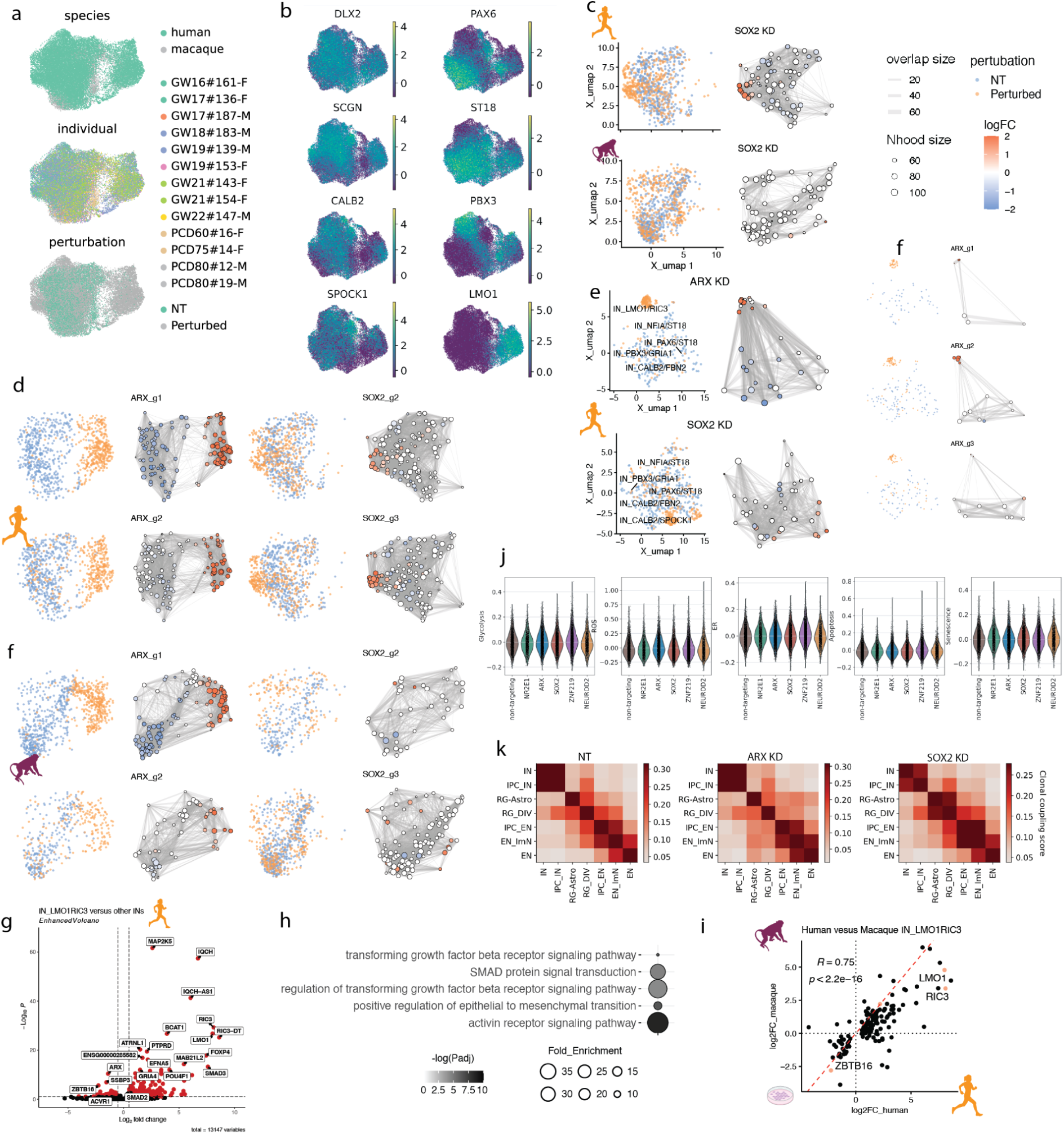
Ectopic IN subtypes in human and macaque. a. Integrated UMAPs of lineage-resolved human and macaque INs, colored by species, individual and perturbation. b. UMAPs of marker gene expression in INs. c. UMAPs for differential abundance testing in IN subtypes using Milo in *SOX2* KD at the gene level in human (top) and macaque (bottom). NT cells were subset based on the number of perturbed cells to balance the cell number between conditions. (left) UMAP of subset NT and perturbed cells, colored by perturbation condition and labeled by IN subcluster. (right) Neighborhood graph of differential abundance testing colored by log_2_FC. d. UMAPs for results of differential abundance testing using Milo for *ARX* (left) and *SOX2* (right) KD at the sgRNA level in human. e. UMAPs for results of differential abundance testing using Milo for *ARX* (top) and *SOX2* (bottom) KD at the gene level in the initial human screen. f. UMAPs of results for differential abundance testing using Milo for *ARX* (left) and *SOX2* (right) KD at the sgRNA level in macaque. g. Volcano plot for DEGs in human IN_LMO1/RIC3 compared to other IN subclusters detected in NT. Genes were colored with a threshold of adjusted p value < 0.01 and |log_2_FC| > 0.5. h. Enrichment plot for GO terms in biological processes enriched in DEGs identified in human IN_LMO1/RIC3 cluster using pathfindR^105^. Color denotes -log_10_(adjusted p value) and dot size denotes fold enrichment in each category. i. Scatterplot of log_2_FC of DEGs identified in IN_LMO1/RIC3 in human and macaque. Pearson correlation coefficients r and p values were labeled. Red dashed line represents Y=X. j. Violin plots of gene expression scores of 5 types of cell stress (glycolysis, ER stress, ROS, apoptosis and senescence) in INs under TF perturbations. Increased cell stress was not observed in INs under *ARX* KD. k. Heatmap for lineage coupling score matrix in major cell types identified in NT (left), *ARX* (middle) and *SOX2* (right) KD in human.

**Extended Data Fig. 9:**
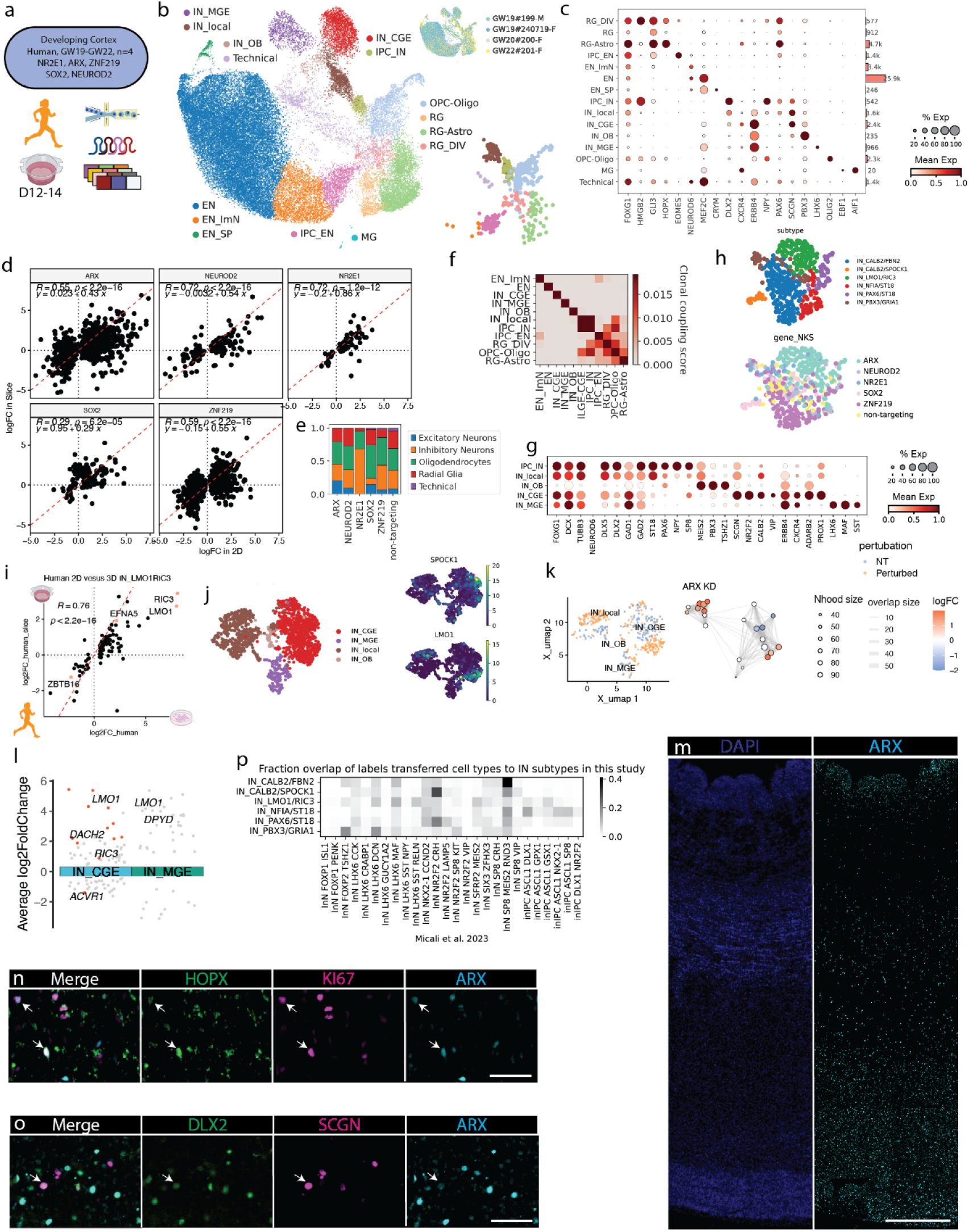
Ectopic IN subtypes in human organotypic slice culture. a. Experimental design for targeted TF perturbation and lineage tracing in organotypic slice culture of 4 human individuals. CRISPRi and STICR lentivirus was locally co-injected to the germinal zone to enrich for RG populations. b. UMAPs showing cell type annotations and individuals (top right), with a UMAP of new born progenies after filtering for cells from multicellular clones on the bottom right. Note that large proportions of postmitotic ENs were labeled and captured but not clonally linked to other cell types. c. Dotplot of the expression of cell type markers for the assigned clusters (left), with a barplot showing numbers of cells detected in the cluster (right). The size of each dot denotes the fractions of cells in the group where the gene is expressed and the color denotes mean gene expression in the group. d. Scatterplot showing log_2_FC in human 2D and slice for DEGs identified in *ARX, NR2E1, SOX2* and *ZNF219* KD. Pearson correlation coefficients and p values were calculated in each TF perturbation and labeled together with regression line equations. Red dashed line represents Y=X. An average Pearson correlation R = 0.6 was observed across perturbations. e. Barplot showing cell class compositions of cells in multicellular clones in each perturbation. f. Heatmap for lineage coupling score matrix in major cell types in the RG lineage. IN_local represents locally derived INs in culture that shares lineage with RG and ENs. g. Dot plot showing markers for different IN subtypes. h. UMAPs of cortical born IN subtypes in IN_local, colored by subtypes and perturbations. i. Scatterplot of log_2_FC of DEGs identified in IN_LMO1/RIC3 in human 2D and slice culture. Pearson correlation coefficients and p values were labeled. Red dashed line represents Y=X. j. UMAPs of all INs identified in slice culture, including post mitotically labeled GE derived clusters (left), and *LMO1* and *SPOCK1* expression (right). k. UMAPs of differential abundance testing in INs using Milo for *ARX* KD. (left) UMAP of NT and perturbed cells, colored by perturbation condition and labeled by IN subcluster. (right) Neighborhood graph of differential abundance testing colored by log_2_FC. l. Volcano plot for DEGs under *ARX* KD in two GE-derived INs defined in slice culture. DEGs with an absolute log_2_FC change more than 0.2 were shown. DEGs with adjusted p value < 0.01 were highlighted in red. m. Immunohistochemistry on GW17 cortex to show expression of ARX, scale bar: 500 μm. n. Immunohiscytochemical labeling of ARX with HOPX and KI67 at subventricular zone, scale bar: 50 μm. o. Immunohiscytochemical labeling of ARX with DLX2 and SCGN at cortical plate, scale bar: 50 μm. p. Heatmap for fraction overlap between IN clusters identified in the human targeted 2D screen and labels transferred from an *in vivo* atlas of the developing macaque telencephalon^106^

**Extended Data Fig. 10:**
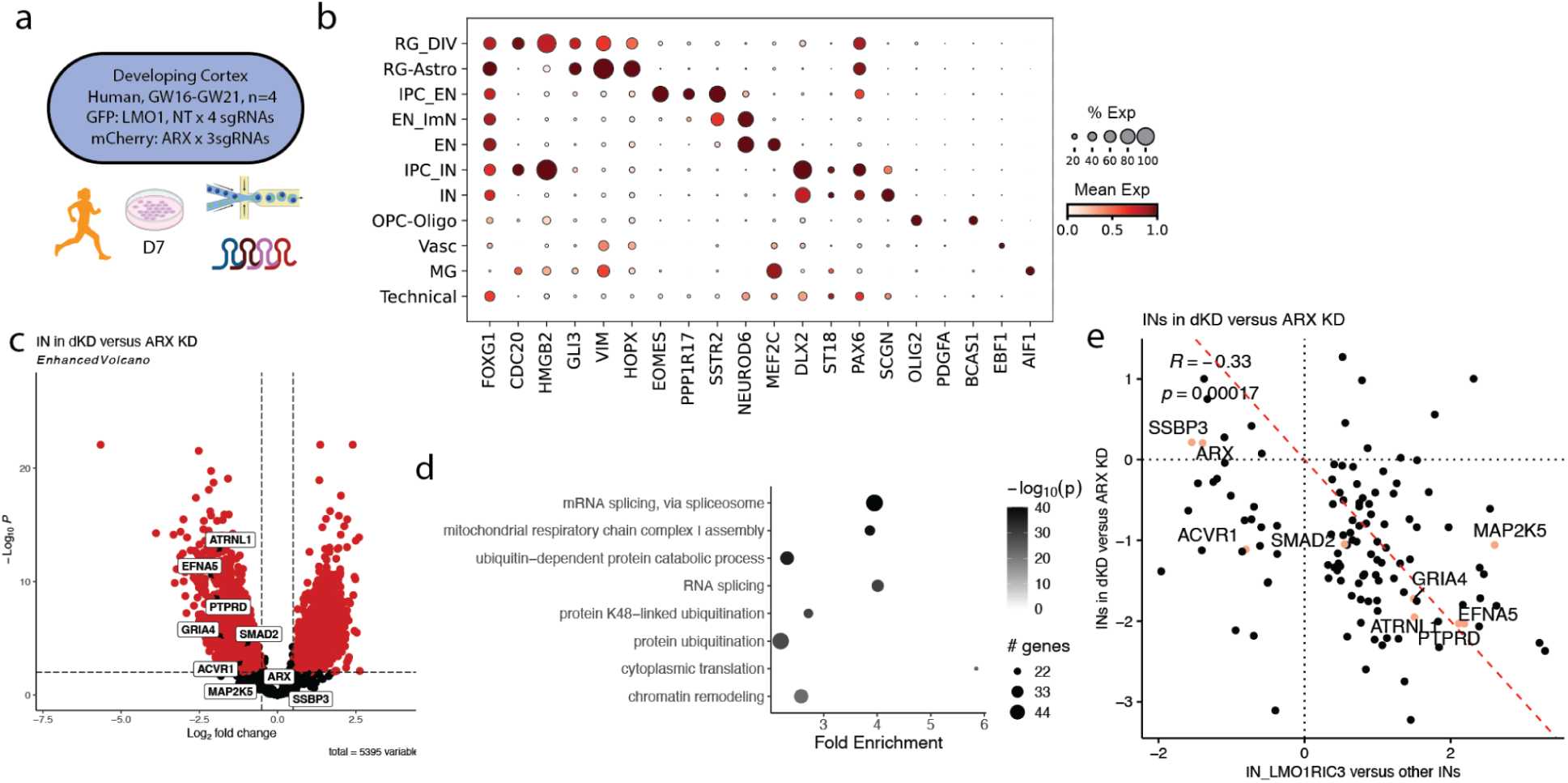
Supplemental analysis for *ARX* and *LMO1* double KD. a. Experimental design for ARX, LMO1 dKD in cortical RG of 4 human individuals. Cells were transduced with GFP expressing library that includes 8 sgRNAs targeting *LMO1* and NT, and mCherry expressing library with 3 sgRNAs targeting *ARX*. GFP and mCherry double positive cells were sorted and captured for scRNA-seq. b. Dotplot of the expression of cell type markers for the assigned clusters (left). c. Volcano plot for DEGs in ARX, LMO1 dKD compared to ARX KD in INs. Genes were colored with a threshold of adjusted p value < 0.01 and |log_2_FC| > 0.5. d. Enrichment plot for GO terms in biological processes enriched in DEGs identified in panel f using pathfindR. Color denotes -log_10_(adjusted p value) and dot size denotes number of DEGs in each category. Enrichment of chromatin related terms show downstream effects of *LMO1* perturbation. e. Scatterplot of log_2_FC of DEGs identified IN_*LMO1/RIC3* in INs (X axis) and log_2_FC in INs of dKD compared to *ARX* KD (Y axis). Significant DEGs (adjusted p value < 0.05 and |log_2_FC| > 0.5) identified in IN_*LMO1/RIC3* from the targeted human screen were shown in this plot. Pearson correlation coefficients and p values were labeled. Red dashed line denotes Y=-X. An overall negative correlation of gene expression changes suggested a rescue of the global transcriptional landscape in dKD.

## Supplementary Table

Supplementary Table 1. Gene regulatory networks inferred from 2 public single cell multiome datasets in the developing human cortex

Supplementary Table 2. Sequence of sgRNAs used in this study

Supplementary Table 3. Batch information for sequencing runs and individual pooling

Supplementary Table 4. KD efficiency of each sgRNA with cell class resolution on human differentiation D7

Supplementary Table 5. Differential expression analysis on human differentiation D7

Supplementary Table 6. Metadata and quantification of cell composition changes using flow cytometry

Supplementary Table 7. Differential expression analysis on macaque differentiation D7

Supplementary Table 8. Differential expression analysis on ectopic IN_LMO1/RIC3 induced by ARX KD

## Methods

### Data and code availability

Raw sequencing data and processed data will be made available through dbGaP upon publication. Any additional information required to reanalyze the data reported in this paper is available from the lead contact upon request.

### Tissue processing and cell culture

#### Tissue samples

De-identified human tissue samples were collected with previous patient consent in strict observance of the legal and institutional ethical regulations. Protocols were approved by the Human Gamete, Embryo, and Stem Cell Research Committee (institutional review board) at the University of California, San Francisco.

The Primate Center at the University of California, Davis, provided 4 specimens of cortical tissue from PCD60 (n=1), PCD75 (*n* = 1) and PCD80 (*n* = 2) macaques. All animal procedures conformed to the requirements of the Animal Welfare Act, and protocols were approved before implementation by the Institutional Animal Care and Use Committee at the University of California, Davis.

#### Cell culture and lentiviral transduction

For cryopreservation, tissue samples were cut into small pieces, placed in Bambanker (NIPPON Genetics, BB02), frozen in CoolCell at -80 and transferred to liquid nitrogen within 2 weeks. For single cell dissociation, each cryovial was thawed in 37°C warm water and placed in a vial containing a pre-warmed solution of Papain (Worthington Biochemical Corporation, LK003153) solution supplemented with 5% Trehalose (Fisher Scientific, BP268710) that was prepared according to the manufacturer protocol for 10 min at 37°C. After approximately 30 min incubation, tissue was triturated following manufacturer protocol. Cells were plated on a 24-well tissue culture dish coated with 0.1% PEI (Sigma, P3143), 5 μg/ml Biolamina LN521 (Invitrogen, 23017-015) and at a density of 500,000 cells/cm^2^. Expansion media contained Insulin (Thermo, A1138IJ) -Transferrin (Invitria, 777TRF029-10G) -Selenium (Sigma, S5261-10G), 1.23 mM ascorbic acid (Fujifilm/Wako, 321-44823), 1% polyvinyl alcohol (PVA) (Sigma, P8136-1KG), 100 μg/ml primocin (Invivogen, ant-pm-05), 20 ng/ml FGF2 (Preprotech, 100-18B), 20 ng/ml EGF (Preprotech, AF100-15) in DMEM-F12 (Corning, MT10092CM), supplemented with ROCK inhibitor CEPT cocktail^107^. For lentiviral transduction, CRISPRi and STICR lentivirus was added to culture media on day 5 at roughly 1:500 and 1:5000 dilution, respectively. After 6 h, the virus-containing medium was removed and replaced with fresh medium. 7 days after infection, expansion media was removed and replaced with differentiation media containing Insulin-Transferrin-Selenium, 1.23 mM ascorbic acid, 1% PVA, 100 μg/ml primocin, 20 ng/ml BDNF (Alomone Labs, B-250) in DMEM-F12. 7 days after differentiation, cultures were dissociated using papain supplemented with 5% Trehalose and GFP positive or GFP and mCherry co-positive cells were isolated by fluorescence-activated cell sorting (FACS) on BD Aria Fusion, resuspended in 0.2% BSA in PBS, and captured with 10X Chromium v3.1 HT kit (PN-1000348) or Illumina Single Cell CRISPR Library kit.

#### Organotypic slice culture and lentiviral transduction

For organotypic slice culture experiments, samples were embedded in 3% low-melting-point agarose (Fisher, BP165-25) and then cut into 300-μm sections perpendicular to the ventricle on a Leica VT1200S vibrating blade microtome in oxygenated artificial cerebrospinal fluid containing 125 mM NaCl, 2.5 mM KCl, 1 mM MgCl2, 1 mM CaCl2 and 1.25 mM NaH2PO4. Slices were cultured slice media containing in Insulin-Transferrin-Selenium, 1.23 mM ascorbic acid, 1% PVA, 100μg/ml primocin, Glutamax (Invitrogen, 35050061), 1 mg/mL BSA, 15 μM Uridine (Sigma, U3003-5G), 1 μg/ml reduced glutathione (Sigma), 1 μg/ml (+)-α-Tocopherol acetate (Sigma, t3001-10g), 0.12 μg/ml Linoleic (Sigma, L1012) and Linolenic acid (Sigma, L2376), 10mg/mL DHA (Cayman, 10006865), 5mg/mL AHA (Cayman, 90010.1), 20ng/ml BDNF (Alomone Labs, B-250) in DMEM-F12. CEPT cocktail was added on the first day. Lentiviral transduction was performed on the following day locally at the germinal zone to preferentially label neural progenitor cells. 24 h after transduction, virus-containing medium was replaced with fresh medium and daily half-medium replacement was performed. 12-14 days after transduction, cultures were dissociated using papain supplemented with 5% Trehalose, and GFP and mCherry co-positive cells were isolated by FACS and captured with 10X Chromium v3.1 HT kit (PN-1000348).

#### Immunocytochemistry

Cells were fixed in 2% PFA for 15 min at room temperature and then in ice cold 90% MeOH for 10 min. After 1 hr incubation in blocking solution (5% BSA, 0.3% Triton-X in PBS), cells were incubated with primary antibodies with the following dilution in blocking solution at room temperature for 1 hr: mouse-EOMES (Thermofisher, 14-4877-82) 1:500, rabbit-NEUROD2 (Abcam, ab104430) 1:500, goat-SOX9 (R&D, AF3075) 1:300, mouse-KI67 (BD, 550609) 1:500, mouse-DLX2 (Santa Cruz Biotechnology, sc-393879) 1:200. The cells were then washed with PBS for 3 times and incubated with secondary antibodies in blocking solution for 1 hr at room temperature, and counterstained with Hoechst 33258. Finally, after PBS washes, digital image acquisition was performed using an Evos M7000 microscope.

#### Flow cytometry

Intracellular staining was performed with Foxp3 transcription factor staining kit (Invitrogen, 00-5523-00) according to the manufacturer protocol. Briefly, cell culture was dissociated with Accutase (STEMCELL Technologies, 07920) supplemented with 5% Trehalose (Fisher Scientific, BP268710) and fixed in Foxp3 fixation buffer for 30 min at room temperature. Cells were then washed twice with Foxp3 permeabilization buffer and then stained with primary antibodies mouse-DLX2 (Santa Cruz Biotechnology, sc-393879) and rabbit-NEUROD2 (Abcam, ab104430) at 1:100 dilution. After washing with Foxp3 permeabilization buffer, cells were stained with secondary antibodies donkey anti-mouse-488 and donkey anti-rabbit-647 (Invitrogen) at 1:200. Finally, cells were stained with conjugated antibodies KI67-421 (BD, 562899), SOX2-PerCP-Cy5.5 (BD, 561506), EOMES-PE-Cy7 (Invitrogen, 25-4877-42) at 1:100, washed twice with Foxp3 permeabilization buffer and resuspended in 0.2% BSA in PBS. Cells were then directly analyzed by flow cytometry. Data was analyzed with FlowJo. Fractions of marker positive populations were normalized to the mean value of 2 NT sgRNAs in each individual and timepoint and paired two-sided Wilcoxon test was performed in each sgRNA compared to both NT sgRNAs to identify significant changes in cell type composition.

#### Immunohistochemistry

Primary cortical tissue were fixed with 4% paraformaldehyde (PFA) in PBS overnight, washed three times with PBS, then placed in 10% and 30% sucrose in PBS overnight, sequentially, and embedded in OCT for sectioning to 20 μm.

All samples were blocked in blocking solution (5% BSA, 0.3% Triton-X in PBS) for 1 hr. Primary and secondary antibodies were diluted in blocking solution. Samples were incubated in primary antibody solution overnight at 4C, then washed three times with PBS at room temperature. Samples were then incubated in a secondary antibody solution with DAPI for 1 hr at room temperature and then washed three times with PBS before mounting samples on slides with Fluoromount (Invitrogen, 0100-20). Primary antibodies included in this analysis include mouse-DLX2 (Santa Cruz Biotechnology, sc-393879), 1:50; rabbit-SCGN (Millipore-sigma, HPA006641), 1:500; rabbit-KI67 (Vector, VP-K451), 1:500; mouse-HOPX (Santa Cruz Biotechnology, sc-398703), 1:50; sheep-ARX (R&D, AF7068SP), 1:500. Secondary antibodies in this study include: donkey anti mouse 647 (1:1000, A31571, Thermo), donkey anti rabbit 555 (1:1000, A11012, Thermo) and donkey anti goat 488 (1:1000, A11055, Thermo). Images were collected using 20x air objectives on an Evos M7000 microscope, and processed using ImageJ/Fiji.

#### Target TF selection through enhancer-driven gene regulatory network (eGRN) inference

Two public datasets of single cell multi omic developing human cortex^10,12^ were used for eGRN inference. The Wang et al. dataset was randomly subset to 50,000 cells. Cells were grouped into meta cells using SEACells^108^ to overcome data sparsity and noise. On average 1 meta cell contains 75 single cells and meta cells representing all the cell types originally presented in the dataset were input to the SCENIC+ workflow. First, MACS2 was used for peak calling in each cell type for each species. A consensus peak set for each species was generated using the TGCA iterative peak filtering approach following the pycisTopic workflow. Next, the merged consensus peaks were summarized into a peak-by-nuclei matrix and topic modeling was performed on the matrix by pycisTopic using default parameters, and the optimal number of topics (15) was determined based on 4 different quality metrics provided by SCENIC+, including log likelihood. We applied three different methods in parallel to identify candidate enhancer regions by selecting regions of interest through (1) binarizing the topics using the Otsu method; (2) taking the top 3,000 regions per topic; (3) calling differentially accessible peaks on the imputed matrix using a Wilcoxon rank sum test (log_2_FC > 1.5 and Benjamini–Hochberg adjusted P values < 0.05). To assess whether the candidate enhancers were linked to a given TF, Pycistarget and discrete element method (DEM) based motif enrichment analysis were implemented. Next, eGRNs, defined as TF-region-gene network consisting of (1) a specific TF, (2) all regions that are enriched for the TF-annotated motif, (3) all genes linked to these regions, were determined by a wrapper function provided by SCENIC+ using the default parameters. eGRNs were then filtered using following criteria: (1) Only eGRNs with more than ten target genes and positive region-to-gene relationships were retained; (2) eGRNs with an extended annotation was only kept if no direct annotation is available; (3) Only genes with top TF-to-gene importance scores (rho > 0.05) were selected as the target genes for each eGRN. Specificity scores were calculated using the RSS algorithm based on region-or gene-based eGRN enrichment scores (AUC scores). To predict potential effects of TF repression on RG differentiation, *in silico* knockdown simulation was applied following the SCENIC+ workflow to a subset of TFs that showed high correlation between TF expression and target region enrichment scores. Briefly, a simulated gene expression matrix was generated by predicting the expression of each gene using the expression of the predictor TFs, while setting the expression of the TF of interest to zero. The simulation was repeated over 5 iterations to predict indirect effects. Cells were then projected onto a eGRN target gene-based PCA embedding and the shift of the cells in the original embedding was estimated based on eGRN gene-based AUC values calculated using the simulated gene expression matrix. Metrics including (1) normalized TF expression in RG, (2) scaled TF motif accessibility and target gene expression in RG, (3) predicted eGRN sizes, and (4) cell-type specificity (RSS) of eGRNs, (5) predicted transcriptomic consequences in the RG lineage were used to select TF targets for this study (Supplementary Table 1).

### Plasmids and lentivirus production

#### Plasmids

The all-in-one CRISPRi plasmid encoding dCas9-KRAB and sgRNA were obtained from Addgene (Addgene, 71237). Capture sequence 1(CS1) was cloned into the vector by Vectorbuilder to enable sgRNA capture with 10X Genomics single cell capture. mCherry-H2B was cloned into the vector to substitute GFP for lineage tracing and flow cytometry experiments. For single sgRNA cloning,the oligonucleotides containing 20bp protospacer and overhang were obtained from Integrated DNA Technologies and cloned into the BsmBI-v2 (New England Biolabs, R0739) digested backbone through T4 ligation (New England Biolabs, M0202). Protospacers were obtained from dual sgRNA CRISPRi libraries^109^ or Dolcetto^51^. For experiments in macaques, protospacers that were uniquely mapped to rhemac10 were selected. STICR plasmids (Addgene #180483, #186334, #186335) used for lineage tracing were obtained from the Nowakowski lab.

#### sgRNA library construction

Oligonucleotide pools were synthesized by Twist Bioscience. BsmBI recognition sites were appended to each sgRNA sequence along with the appropriate overhang sequences for cloning into the sgRNA expression plasmids, as well as primer sites to allow differential amplification of subsets from the same synthesis pool. The final oligonucleotide sequence was: 5′-[Forward Primer]CGTCTCA*CACCG*[sgRNA, 20 nt]*GTTT*CGAGACG[Reverse Primer].

Primers were used to amplify individual subpools using 50 μL 2x NEBNext Ultra II Q5 Master Mix (New England Biolabs, M0544S), 20 μL of oligonucleotide pool (∼20 ng), 0.5 μL of primer mix at a final concentration of 0.5 μM, and 29 μL nuclease-free water. PCR cycling conditions: (1) 98°C for 30 s; (2) 98°C for 10 s; (3) 68°C for 30 s; (4) 72°C for 30 s; (5) go to (2), x 8; (5) 72°C for 2 min.

The resulting amplicons were PCR-purified (Zymo, D4060) and cloned into the library vector via Golden Gate cloning with Esp3I (Thermo Scientific, ER0451) and T7 ligase (New England Biolabs, M0318); the library vector was pre-digested with BsmBI-v2 (New England Biolabs, R0739). The ligation product was electroporated into MegaX DH10B T1R Electrocomp Cells (Invitrogen, C640003) and grown at 30°C for 24 h in 200 mL LB broth with 100 μg/mL carbenicillin. Library diversity and sgRNA representation were assessed through PCR amplicon sequencing.

#### Lentivirus production

Lentivirus was produced in HEK293T cells. HEK293Ts were seeded at a density of 80,000 cells/cm^2^ 24 hr prior to transfection. Transfection was performed using Lipofectamine 3000 (Invitrogen, L3000001) transfection reagent according to the manufacturer’s protocol. 18 hr post-transfection, the media was replaced and supplemented with 1X ViralBoost (Alstem, VB100). Supernatant was collected at 48 hr post-transfection and concentrated roughly at 1:100 with lentivirus precipitation solution (Alstem, VC100).

### Generation and analysis of scRNA-seq libraries

#### scRNA-seq library generation

The manufacturer-provided protocol (CG000421 Rev D) was used to generate 10x single cell gene expression and sgRNA libraries. Samples from multiple individuals were pooled (Supplementary Table 3) and each 10X lane was loaded around 100,000 cells in total. sgRNA libraries were separately amplified from each 10X cDNA library using 10X 3’ Feature Barcode Kit (PN-1000262). To generate STICR barcode libraries, 10 μl of 10X cDNA library was used as template in a 50 μl PCR reaction containing 25 μl Q5 Hot Start High Fidelity 2X master mix (NEB, M0494) and STICR barcode read 1 and 2 primers (0.5 μM, each) described in Delgado et al.^1^ using the following program: 1, 98 °C, 30 s; 2, 98 °C, 10 s; 3, 62 °C, 20 s; 4, 72 °C, 10 s; 5, repeat steps 2–4 15 times; 6, 72 °C, 2 min; 7, 4 °C, hold. Following PCR amplification, a 0.8–0.6 dual-sided size selection was performed using SPRIselect Bead (Beckman Coulter, B23318).

For *ARX* and *LMO1* double KD (dKD) experiments, cells are captured with Illumina Single Cell CRISPR Library kit where direct sgRNA capture was enabled. Libraries were prepared following the manufacturer-provided protocol (FB0004762; FB0002130).

#### Alignments and quality control

Libraries were sequenced on Illumina NovaSeq platforms to the depth of roughly 20,000-30,000 reads/cell for transcriptomics and 5000 reads/cell for sgRNA and STICR libraries.

10X transcriptomic libraries together with sgRNA and STICR libraries were aligned to the hg38 genome with feature barcode reference (Supplementary Table 2) using CellRanger-7.2.0. Aligned cell/transcript counts were then processed with Cellbender^110^ in order to identify and remove background reads. The resulting counts were processed by Scanpy^111^ to remove cells containing fewer than 1000 genes, a high abundance of mitochondrial reads (greater than 15% of total transcripts), or a high abundance of ribosomal reads (greater than 40% of total transcripts). Illumina Single Cell CRISPR libraries were aligned using DRAGEN Single Cell RNA v4.4.5 and cells with fewer than 500 genes or high abundance of mitochondrial or ribosomal reads were removed.

For runs with pooled humans and macaques (Supplementary Table 3), libraries were aligned to a chimaeric hg38/rheMac10 genome using CellRanger in order to identify cross-species multiplets; cells identified as such were removed from downstream analyses.

#### Demultiplexing of individuals from pooled sequencing and doublet removal

CellSNP-lite followed by Vireo^112,113^ were used to identify different individuals based on reference-free genotyping using candidate SNPs identified from1000 Genome Project. Sex information was acquired through PCR-based genotyping using genomic DNA extracted from each individual. Each individual was assigned based on sex and pooling information (Supplementary Table 3). Same individuals from different 10X lanes were merged based on identical SNP genotypes (Extended Data Fig. 1c). Interindividual doublets and unassigned cells were removed from downstream analyses.

#### sgRNA and lineage barcode assignments

Cellbouncer(https://github.com/nkschaefer/cellbouncer) was used for sgRNA assignment using the sgRNA count matrix obtained from Cellranger. ‘Effective sgRNA’ was defined for each cell based on assigned sgRNAs after collapsing NT sgRNA. For example, cells assigned with NT sgRNA and SOX2-targeting sgRNA are classified as ‘Effective sgRNA’=SOX2. ‘Target Gene’ was defined for each cell based after collapsing NT sgRNA and sgRNAs sharing the same target genes. For example, cells infected with 2 different SOX2-targeting sgRNAs are classified as ‘Target Gene’=SOX2 and included for analyzing SOX2 KD phenotypes at the gene level. Multi-infected cells with sgRNAs targeting different genes were assigned to have more than 1 ‘Target Gene’ and removed from downstream analyses.

KD efficiency was calculated in each cell class following the DEseq2 workflow, where single cell data was pseudo-bulked by cell class and sgRNAs and different individuals were used as replicates. We reasoned that KD efficiency can vary between cell types and classes depending on the baseline expression level of the target gene. The lowest log2FC among cell classes was used for downstream filtering. sgRNAs that had maximum KD less than 25% (log2FC > -0.4, 18 sgRNAs) were removed from downstream analyses.

STICR barcodes for lineage tracing experiments were aligned and assigned using a modified NextClone^114^ workflow that allows for whitelisting. The pipeline is available at: https://github.com/cnk113/NextClone. Individual barcodes were filtered by at least 3 reads supporting a single UMI and at least 2 UMI to call cells with a barcode. Clone calling was done using CloneDetective^114^.

#### Comparison to public datasets and cell type annotation

The developing human cortex multiomic dataset, including metadata and aligned cellranger output, was obtained from Wang et al.^12^ and used for reference mapping. The reference model was built with scvi-tools^115^ using top 2500 variable genes and label transfer to the *in vitro* query dataset generated in this study was performed to examine correspondence of cell type assignment based on the *in vivo* dataset. Cell type annotation was then performed based on marker expression as well as predictions from scANVI.

The initial human screen generated in this study was annotated and used as a reference dataset to map cell populations collected from from lineage-resolved targeted screen in human and macaque and the ARX, LMO1 dKD experiment to minimize the impacts of batch effects and species differences on cell type annotation.

Datasets from public organoid^53^, iNeuron^102^ and FeBO^52^ studies were used to compare fidelity of *in vitro* specified cell types between different systems. Reference mapping to the abovementioned *in vivo* dataset and label transfer were performed to examine correspondence of cell type assignment based on the *in vivo* dataset. Pearson correlation coefficients between datasets were calculated using top 25 markers in each cell type identified from the *in vivo* dataset to show the fidelity of cell identities in this study. To assess the level of cellular stress, gene scores for glycolysis, ER stress, ROS, apoptosis and senescence were calculated using the score_genes function in Scanpy using gene sets obtained from the MSigDB database^116,117^. NT control cells of all cell types were subset from data obtained in this study for this comparison. Each dataset was then randomly downsampled to 5,000 cells to ensure balanced representation between datasets.

#### Trajectory analysis using pseudotime and RNA velocity

The initial human screen data was subset to NT cells to remove effects from TF perturbation on inferring pseudotime and RNA velocity. Pseudotime was calculated using scFates^118^ by tree learning with SimplePPT and setting the root node within the radial glia class on ForceAtlas2 embedding. Excitatory and inhibitory trajectories were defined as major branches of the principal graph that led to distinct sets of clusters. The Velocyto^119^ pipeline to quantify spliced and unspliced reads from cellranger output. Highly variable genes were separately calculated using spliced and unspliced matrices and top 3000 genes were used for inferring RNA velocity using scVelo^120^.

#### Differential composition and gene expression analysis

Cells containing sgRNAs targeting single genes were subset and sgRNAs targeting the same gene were aggregated for downstream analyses at the gene level. Composition changes in each cluster under each TF perturbation were quantified using DCATS^104^, which detects differential abundance using a beta-binomial generalized linear model (GLM) model and returns the estimated coefficients and p-values. To detect compositional changes in a finer resolution, Milo^101^ was used to quantify differential abundance in a label-free manner. Briefly, perturbed and NT cells were subset to ensure the balance of total cell numbers between the 2 groups. Neighborhoods were constructed based on KNN graphs, and then cell abundances under each condition in each neighborhood were tested against NT using design = ∼ stage + sex + batch + perturbation.

Cluster-aware differential gene expression analysis was performed following the DEseq2^54^ pipeline using NT cells as reference. When pseudobulking within cell types, conditions that have less than 5 cells per cell type or less than 2 biological replicates were removed from downstream analysis. DEGs identified under more than one perturbation across 44 TFs perturbations in the initial human screen were defined as ‘convergent DEGs’ and GO enrichment analysis was performed using non-convergent DEGs as background. Fisher’s exact test was performed to identify significant overlap between convergent DEGs and 7 sets of disease-associated genes (See below) using non-convergent DEGs as background.

To prioritize TFs whose repression led to strongest cellular and transcriptional and consequences in the initial human screen, absolute values of estimated coefficient from DCATS and numbers of DEGs from DEseq2 across 7 cell types in the EN and IN lineage were summed to quantify accumulated effects of TF repression in composition and gene expression, respectively. To test the robustness of prioritized TFs, scCoda^103^ was applied to calculate log fold change in cell abundance. Euclidean and energy distance between each TF perturbation were calculated with Pertpy^121^ to compare the global transcriptional landscape post perturbation.

#### Neurological disorder-associated genes from previous studies

Genes significantly associated with neurodevelopmental and neurological disorders were obtained from:

- ASD: SFARI gene database^83^, score 1
- NDD: Fu et al.^84^, Supplementary Table 11
- MDD: Howard et al.^122^, Supplementary Table 9
- SCZ: Trubetskoy et al.^123^, Supplementary Table 12
- BP: Mullins et al.^124^, Supplementary Table 4
- ADHD: Demontis et al.^125^, Supplementary Table 7
- AD: Bellenguez et al.^126^, Supplementary Table 5

#### Lineage coupling and clonal clustering

Clones with less than 3 cells and clones that have conflicted sgRNA assignments were removed from the clonal analysis. One human individual (GW17#187-M) with overall low cellular coverage was dropped from the following analyses. Cospar^86^ was used to calculate the fate coupling scores, defined as the normalized barcode covariance between different cell types.

scLiTr^87^ was used to identify and cluster fate biased clones by training a neural network to predict clonal labels of nearest neighbours for each clonally labelled cell. To minimize batch effects in cluster identification, we used integrated lineage-resolved data from both human and macaque to define 5 clusters with distinct fate biases. The obtained results were exported and a paired 2-sided Wilcoxon test was performed comparing the mean fractions of clonal clusters between NT and each TF perturbation. Wilcoxon rank-sum test through Scanpy was used to identify marker genes in the radial glia class (RG-Astro and RG_DIV) between different fate clusters in human. Monocle3^127^ was used to construct pseudotime trajectory at the clonal level and using the scLiTr output separately in each species. The median pseudotime in each individual was calculated under each perturbation and used for paired two-sided Wilcoxon tests to evaluate the changes on clonal pseudotime. To fit gene expression along the clonal pseudotime, the radial glia class from multicellular clones was subset and assigned with pseudotime values.

#### IN subtype identification and differential gene expression

To identify IN subtypes, the IN cluster from the initial human screen was subset and re-clustered into 6 subtypes, each expressing distinct markers identified through Scanpy using Wilcoxon rank-sum test. Reference mapping and label transfer were then performed using lineage-resolved human and macaque INs as query dataset to identify equivalent clusters. For cell abundance testing in IN subtypes in each perturbation, NT and perturbed INs were subset and the same Milo^101^ pipeline described above was applied to identify differential abundance within the IN cluster. Differential expression analysis was performed using DEseq2 contrasting the IN_LMO1/RIC3 cluster with other physiological subtypes identified in unperturbed INs (IN_NFIA/ST18, IN_PAX6/ST18, IN_CALB2/FBN2, IN_PBX3/GRIA1) in each species. GO terms enrichment in biological processes in human IN_LMO1/RIC3 cluster were identified using pathfindR^105^.

#### *ARX* and *LMO1* double KD (dKD)

GFP expressing CRISPRi library targeting *LMO1* and NT controls, together with mCherry expressing vector targeting *ARX* were used to achieve dKD and double infected cells expressing both colors were sorted and captured with Illumina Single Cell CRISPR Library Kit. The resulting library was aligned with DRAGEN Single Cell RNA v4.4.5 and reference mapped to the initial human screen data. sgRNA assignment was done using cellbouncer and double infected cells (*ARX, LMO1* dKD or *ARX*, NT KD) were integrated with previous data from the targeted human screen for downstream analysis. Differential gene expression was performed using DEseq2 by contrasting *ARX, LMO1* dKD against *ARX* KD. Cluster-free differential abundance testing was performed using Milo after integrating with previous batches to compare NT, ARX KD and dKD.

## Notes

### Competing Interest Statement

The authors have declared no competing interest.

## References

1. Delgado, R. N. et al. Individual human cortical progenitors can produce excitatory and inhibitory neurons. Nature 601, 397–403 (2021).

2. Lui, J. H., Hansen, D. V. & Kriegstein, A. R. Development and Evolution of the Human Neocortex. Cell vol. 146 332 Preprint at 10.1016/j.cell.2011.07.005 (2011).

3. Chen, B. et al. The Fezf2-Ctip2 genetic pathway regulates the fate choice of subcortical projection neurons in the developing cerebral cortex. Proc. Natl. Acad. Sci. U. S. A. 105, 11382–11387 (2008).

4. Arlotta, P., Molyneaux, B. J., Jabaudon, D., Yoshida, Y. & Macklis, J. D. Ctip2 controls the differentiation of medium spiny neurons and the establishment of the cellular architecture of the striatum. J. Neurosci. 28, 622–632 (2008).

5. Britanova, O. et al. Satb2 is a postmitotic determinant for upper-layer neuron specification in the neocortex. Neuron 57, 378–392 (2008).

6. Lai, T. et al. SOX5 controls the sequential generation of distinct corticofugal neuron subtypes. Neuron 57, 232–247 (2008).

7. Azim, E., Jabaudon, D., Fame, R. M. & Macklis, J. D. SOX6 controls dorsal progenitor identity and interneuron diversity during neocortical development. Nat. Neurosci. 12, 1238–1247 (2009).

8. Telley, L. et al. Sequential transcriptional waves direct the differentiation of newborn neurons in the mouse neocortex. Science 351, 1443–1446 (2016).

9. Ziffra, R. S. et al. Single-cell epigenomics reveals mechanisms of human cortical development. Nature 598, 205–213 (2021).

10. Trevino, A. E. et al. Chromatin and gene-regulatory dynamics of the developing human cerebral cortex at single-cell resolution. Cell 184, 5053–5069.e23 (2021).

11. Mannens, C. C. A. et al. Chromatin accessibility during human first-trimester neurodevelopment. Nature 1–8 (2024).

12. Wang, L. et al. Molecular and cellular dynamics of the developing human neocortex. Nature 1–10 (2025).

13. Dixit, A. et al. Perturb-Seq: Dissecting Molecular Circuits with Scalable Single-Cell RNA Profiling of Pooled Genetic Screens. Cell 167, 1853–1866.e17 (2016).

14. Adamson, B. et al. A Multiplexed Single-Cell CRISPR Screening Platform Enables Systematic Dissection of the Unfolded Protein Response. Cell 167, 1867–1882.e21 (2016).

15. Otani, T., Marchetto, M. C., Gage, F. H., Simons, B. D. & Livesey, F. J. 2D and 3D Stem Cell Models of Primate Cortical Development Identify Species-Specific Differences in Progenitor Behavior Contributing to Brain Size. Cell Stem Cell 18, 467–480 (2016).

16. Pollen, A. A. et al. Molecular identity of human outer radial glia during cortical development. Cell 163, 55–67 (2015).

17. Geschwind, D. H. & Rakic, P. Cortical evolution: judge the brain by its cover. Neuron 80, 633–647 (2013).

18. Lindhout, F. W., Krienen, F. M., Pollard, K. S. & Lancaster, M. A. A molecular and cellular perspective on human brain evolution and tempo. Nature 630, 596–608 (2024).

19. Zhou, Y., Song, H. & Ming, G.-L. Genetics of human brain development. Nat. Rev. Genet.25, 26–45 (2024).

20. Libé-Philippot, B. & Vanderhaeghen, P. Cellular and molecular mechanisms linking human cortical development and evolution. Annu. Rev. Genet. 55, 555–581 (2021).

21. Marín, O. & Rubenstein, J. L. A long, remarkable journey: tangential migration in the telencephalon. Nat. Rev. Neurosci. 2, 780–790 (2001).

22. Hansen, D. V. et al. Non-epithelial stem cells and cortical interneuron production in the human ganglionic eminences. Nat. Neurosci. 16, 1576–1587 (2013).

23. Ma, T. et al. Subcortical origins of human and monkey neocortical interneurons. Nat. Neurosci. 16, 1588–1597 (2013).

24. Bandler, R. C. et al. Single-cell delineation of lineage and genetic identity in the mouse brain. Nature 601, 404–409 (2022).

25. Kim, S. N. et al. Cell lineage analysis with somatic mutations reveals late divergence of neuronal cell types and cortical areas in human cerebral cortex. bioRxiv (2023) doi:10.1101/2023.11.06.565899.

26. Chung, C. et al. Cell-type-resolved mosaicism reveals clonal dynamics of the human forebrain. Nature 629, 384–392 (2024).

27. Wang, L. et al. Molecular and cellular dynamics of the developing human neocortex. Nature 1–10 (2025).

28. Li, C. et al. Single-cell brain organoid screening identifies developmental defects in autism. Nature 621, 373–380 (2023).

29. Fleck, J. S. et al. Inferring and perturbing cell fate regulomes in human brain organoids. Nature 1–8 (2022).

30. Gordon, A. et al. Developmental convergence and divergence in human stem cell models of autism spectrum disorder. Neuroscience (2024).

31. Jin, X. et al. In vivo Perturb-Seq reveals neuronal and glial abnormalities associated with autism risk genes. Science 370, eaaz6063 (2020).

32. Zheng, X. et al. Massively parallel in vivo Perturb-seq reveals cell-type-specific transcriptional networks in cortical development. Cell 187, 3236–3248.e21 (2024).

33. Sherman, M. H., Bassing, C. H. & Teitell, M. A. Regulation of cell differentiation by the DNA damage response. Trends Cell Biol. 21, 312–319 (2011).

34. Friskes, A. et al. Double-strand break toxicity is chromatin context independent. Nucleic Acids Res. 50, 9930–9947 (2022).

35. Merkle, F. T. et al. Human pluripotent stem cells recurrently acquire and expand dominant negative P53 mutations. Nature 545, 229–233 (2017).

36. Kilpinen, H. et al. Common genetic variation drives molecular heterogeneity in human iPSCs. Nature 546, 370–375 (2017).

37. Strano, A., Tuck, E., Stubbs, V. E. & Livesey, F. J. Variable outcomes in neural differentiation of human PSCs arise from intrinsic differences in developmental signaling pathways. Cell Rep. 31, 107732 (2020).

38. Bhaduri, A. et al. Cell stress in cortical organoids impairs molecular subtype specification. Nature 578, 142–148 (2020).

39. Gilbert, L. A. et al. Genome-scale CRISPR-mediated control of gene repression and activation. Cell 159, 647–661 (2014).

40. Gilbert, L. A. et al. CRISPR-mediated modular RNA-guided regulation of transcription in eukaryotes. Cell 154, 442–451 (2013).

41. Stein, J. L. et al. A quantitative framework to evaluate modeling of cortical development by neural stem cells. Neuron 83, 69–86 (2014).

42. Onorati, M. et al. Zika Virus Disrupts Phospho-TBK1 Localization and Mitosis in Human Neuroepithelial Stem Cells and Radial Glia. Cell Rep. 16, 2576–2592 (2016).

43. Pollen, A. A. et al. Establishing Cerebral Organoids as Models of Human-Specific Brain Evolution. Cell 176, 743–756.e17 (2019).

44. Arnold, S. J. et al. The T-box transcription factor Eomes/Tbr2 regulates neurogenesis in the cortical subventricular zone. Genes Dev. 22, 2479–2484 (2008).

45. Mihalas, A. B. & Hevner, R. F. Control of neuronal development by T-box genes in the brain. Curr. Top. Dev. Biol. 122, 279–312 (2017).

46. Ypsilanti, A. R. & Rubenstein, J. L. R. Transcriptional and epigenetic mechanisms of early cortical development: An examination of how Pax6 coordinates cortical development: Mechanisms of Early Cortical Development. J. Comp. Neurol. 524, 609–629 (2016).

47. Manuel, M. N., Mi, D., Mason, J. O. & Price, D. J. Regulation of cerebral cortical neurogenesis by the Pax6 transcription factor. Front. Cell. Neurosci. 9, 70 (2015).

48. Thakore, P. I. et al. Highly specific epigenome editing by CRISPR-Cas9 repressors for silencing of distal regulatory elements. Nat. Methods 12, 1143–1149 (2015).

49. Replogle, J. M. et al. Combinatorial single-cell CRISPR screens by direct guide RNA capture and targeted sequencing. Nat. Biotechnol. 38, 954–961 (2020).

50. Horlbeck, M. A. et al. Compact and highly active next-generation libraries for CRISPR-mediated gene repression and activation. Elife 5, (2016).

51. Sanson, K. R. et al. Optimized libraries for CRISPR-Cas9 genetic screens with multiple modalities. Nat. Commun. 9, 5416 (2018).

52. Hendriks, D. et al. Human fetal brain self-organizes into long-term expanding organoids. Cell 187, 712–732.e38 (2024).

53. He, Z. et al. An integrated transcriptomic cell atlas of human neural organoids. Nature 635, 690–698 (2024).

54. Love, M. I., Huber, W. & Anders, S. Moderated estimation of fold change and dispersion for RNA-seq data with DESeq2. Genome Biol. 15, 550 (2014).

55. Replogle, J. M. et al. Mapping information-rich genotype-phenotype landscapes with genome-scale Perturb-seq. Cell 185, 2559–2575.e28 (2022).

56. Kitamura, K. et al. Mutation of ARX causes abnormal development of forebrain and testes in mice and X-linked lissencephaly with abnormal genitalia in humans. Nat. Genet. 32, 359–369 (2002).

57. Strømme, P., Mangelsdorf, M. E., Scheffer, I. E. & Gécz, J. Infantile spasms, dystonia, and other X-linked phenotypes caused by mutations in Aristaless related homeobox gene, ARX. Brain Dev. 24, 266–268 (2002).

58. Kato, M., Das, S., Petras, K., Sawaishi, Y. & Dobyns, W. B. Polyalanine expansion of ARX associated with cryptogenic West syndrome. Neurology 61, 267–276 (2003).

59. Strømme, P. et al. Mutations in the human ortholog of Aristaless cause X-linked mental retardation and epilepsy. Nat. Genet. 30, 441–445 (2002).

60. Bienvenu, T. et al. ARX, a novel Prd-class-homeobox gene highly expressed in the telencephalon, is mutated in X-linked mental retardation. Hum. Mol. Genet. 11, 981–991 (2002).

61. Wang, Y.-Y., Hsu, S.-H., Tsai, H.-Y. & Cheng, M.-C. Genetic analysis of the NR2E1 gene as a candidate gene of schizophrenia. Psychiatry Res. 293, 113386 (2020).

62. Dennert, N. et al. De novo microdeletions and point mutations affecting SOX2 in three individuals with intellectual disability but without major eye malformations. Am. J. Med. Genet. A 173, 435–443 (2017).

63. Sisodiya, S. M. et al. Role of SOX2 mutations in human hippocampal malformations and epilepsy. Epilepsia 47, 534–542 (2006).

64. Gregor, A. et al. De novo mutations in the genome organizer CTCF cause intellectual disability. Am. J. Hum. Genet. 93, 124–131 (2013).

65. Chen, F. et al. Three additional de novo CTCF mutations in Chinese patients help to define an emerging neurodevelopmental disorder. Am. J. Med. Genet. C Semin. Med. Genet. 181, 218–225 (2019).

66. Runge, K. et al. Disruption of NEUROD2 causes a neurodevelopmental syndrome with autistic features via cell-autonomous defects in forebrain glutamatergic neurons. Mol. Psychiatry 26, 6125–6148 (2021).

67. Sega, A. G. et al. De novo pathogenic variants in neuronal differentiation factor 2 (NEUROD2) cause a form of early infantile epileptic encephalopathy. J. Med. Genet. 56, 113–122 (2019).

68. Kim, H.-G. et al. Disruption of PHF21A causes syndromic intellectual disability with craniofacial anomalies, epilepsy, hypotonia, and neurobehavioral problems including autism. Mol. Autism 10, 35 (2019).

69. Moen, M. J. et al. An interaction network of mental disorder proteins in neural stem cells. Transl. Psychiatry 7, e1082 (2017).

70. Shi, Y. et al. Expression and function of orphan nuclear receptor TLX in adult neural stem cells. Nature 427, 78–83 (2004).

71. Fulp, C. T. et al. Identification of Arx transcriptional targets in the developing basal forebrain. Hum. Mol. Genet. 17, 3740–3760 (2008).

72. Cortés, B. I. et al. Loss of protein tyrosine phosphatase receptor delta PTPRD increases the number of cortical neurons, impairs synaptic function and induces autistic-like behaviors in adult mice. Biol. Res. 57, 40 (2024).

73. Tomita, H. et al. The Protein Tyrosine Phosphatase Receptor Delta Regulates Developmental Neurogenesis. Cell Rep. 30, 215–228.e5 (2020).

74. Yasumura, M. et al. IL1RAPL1 knockout mice show spine density decrease, learning deficiency, hyperactivity and reduced anxiety-like behaviours. Sci. Rep. 4, 6613 (2014).

75. Bani-Yaghoub, M. et al. Role of Sox2 in the development of the mouse neocortex. Dev. Biol. 295, 52–66 (2006).

76. Roy, K. et al. The Tlx gene regulates the timing of neurogenesis in the cortex. J. Neurosci. 24, 8333–8345 (2004).

77. Li, S., Sun, G., Murai, K., Ye, P. & Shi, Y. Characterization of TLX expression in neural stem cells and progenitor cells in adult brains. PLoS One 7, e43324 (2012).

78. Lim, Y., et al. ARX regulates cortical interneuron differentiation and migration. bioRxiv (2024) doi:10.1101/2024.01.31.578282.

79. Chung, C., Girgiss, J. & Gleeson, J. G. A comparative view of human and mouse telencephalon inhibitory neuron development. Development 152, dev204306 (2025).

80. Colombo, E. et al. Inactivation of Arx, the murine ortholog of the X-linked lissencephaly with ambiguous genitalia gene, leads to severe disorganization of the ventral telencephalon with impaired neuronal migration and differentiation. J. Neurosci. 27, 4786–4798 (2007).

81. Takigawa, Y. et al. The transcription factor Znf219 regulates chondrocyte differentiation by assembling a transcription factory with Sox9. J. Cell Sci. 123, 3780–3788 (2010).

82. Nagai, M. et al. Neuronal splicing of the unmethylated histone H3K4 reader, PHF21A, prevents excessive synaptogenesis. J. Biol. Chem. 300, 107881 (2024).

83. Abrahams, B. S. et al. SFARI Gene 2.0: a community-driven knowledgebase for the autism spectrum disorders (ASDs). Mol. Autism 4, 36 (2013).

84. Fu, J. M. et al. Rare coding variation provides insight into the genetic architecture and phenotypic context of autism. Nat. Genet. 54, 1320–1331 (2022).

85. Weinreb, C. & Klein, A. M. Lineage reconstruction from clonal correlations. Proc. Natl. Acad. Sci. U. S. A. 117, 17041–17048 (2020).

86. Wang, S.-W., Herriges, M. J., Hurley, K., Kotton, D. N. & Klein, A. M. CoSpar identifies early cell fate biases from single-cell transcriptomic and lineage information. Nat. Biotechnol. 40, 1066–1074 (2022).

87. Erickson, A. G., et al. Unbiased profiling of multipotency landscapes reveals spatial modulators of clonal fate biases. bioRxiv (2024) doi:10.1101/2024.11.15.623687.

88. Zhang, Y. et al. Cortical neural stem cell lineage progression is regulated by extrinsic signaling molecule Sonic hedgehog. Cell Rep. 30, 4490–4504.e4 (2020).

89. Yang, L., Li, Z., Liu, G., Li, X. & Yang, Z. Developmental origins of human cortical oligodendrocytes and astrocytes. Neurosci. Bull. 38, 47–68 (2022).

90. Li, X. et al. Decoding cortical glial cell development. Neurosci. Bull. 37, 440–460 (2021).

91. Petanjek, Z., Berger, B. & Esclapez, M. Origins of cortical GABAergic neurons in the cynomolgus monkey. Cereb. Cortex 19, 249–262 (2009).

92. Letinic, K., Zoncu, R. & Rakic, P. Origin of GABAergic neurons in the human neocortex. Nature 417, 645–649 (2002).

93. Clowry, G. J. An enhanced role and expanded developmental origins for gamma-aminobutyric acidergic interneurons in the human cerebral cortex. J. Anat. 227, 384–393 (2015).

94. Cho, G., Nasrallah, M. P., Lim, Y. & Golden, J. A. Distinct DNA binding and transcriptional repression characteristics related to different ARX mutations. Neurogenetics 13, 23–29 (2012).

95. Lee, K., Mattiske, T., Kitamura, K., Gecz, J. & Shoubridge, C. Reduced polyalanine-expanded Arx mutant protein in developing mouse subpallium alters Lmo1 transcriptional regulation. Hum. Mol. Genet. 23, 1084–1094 (2014).

96. Jourdon, A. et al. Enhancer-driven regulatory network of forebrain human development provides insights into autism. Genomics (2023).

97. Nowakowski, T. J. et al. Spatiotemporal gene expression trajectories reveal developmental hierarchies of the human cortex. Science 358, 1318–1323 (2017).

98. Zhu, S. et al. LMO1 synergizes with MYCN to promote neuroblastoma initiation and metastasis. Cancer Cell 32, 310–323.e5 (2017).

99. Gao, L. et al. LMO1 plays an oncogenic role in human glioma associated with NF-kB pathway. Front. Oncol. 12, 770299 (2022).

100. Colasante, G. et al. Arx acts as a regional key selector gene in the ventral telencephalon mainly through its transcriptional repression activity. Dev. Biol. 334, 59–71 (2009).

101. Dann, E., Henderson, N. C., Teichmann, S. A., Morgan, M. D. & Marioni, J. C. Differential abundance testing on single-cell data using k-nearest neighbor graphs. Nature Biotechnology 40, 245–253 (2021).

102. Lin, H.-C. et al. NGN2 induces diverse neuron types from human pluripotency. Stem Cell Reports 16, 2118–2127 (2021).

103. Büttner, M., Ostner, J., Müller, C. L., Theis, F. J. & Schubert, B. scCODA is a Bayesian model for compositional single-cell data analysis. Nature Communications 12, 1–10 (2021).

104. Lin, X., Chau, C., Ma, K., Huang, Y. & Ho, J. W. K. DCATS: differential composition analysis for flexible single-cell experimental designs. Genome Biol. 24, 151 (2023).

105. Ulgen, E., Ozisik, O. & Sezerman, O. U. PathfindR: An R package for comprehensive identification of enriched pathways in omics data through active subnetworks. Front. Genet. 10, (2019).

106. Micali, N. et al. Molecular programs of regional specification and neural stem cell fate progression in developing macaque telencephalon. bioRxiv 2022.10.18.512724 (2022) doi:10.1101/2022.10.18.512724.

107. Chen, Y. et al. A versatile polypharmacology platform promotes cytoprotection and viability of human pluripotent and differentiated cells. Nat. Methods 18, 528–541 (2021).

108. Persad, S. et al. SEACells infers transcriptional and epigenomic cellular states from single-cell genomics data. Nat. Biotechnol. 41, 1746–1757 (2023).

109. Replogle, J. M. et al. Maximizing CRISPRi efficacy and accessibility with dual-sgRNA libraries and optimal effectors. Elife 11, e81856 (2022).

110. Fleming, S. J. et al. Unsupervised removal of systematic background noise from droplet-based single-cell experiments using CellBender. Nat. Methods 20, 1323–1335 (2023).

111. Wolf, F. A., Angerer, P. & Theis, F. J. SCANPY: large-scale single-cell gene expression data analysis. Genome Biol. 19, 15 (2018).

112. Huang, Y., McCarthy, D. J. & Stegle, O. Vireo: Bayesian demultiplexing of pooled single-cell RNA-seq data without genotype reference. Genome Biol. 20, 273 (2019).

113. Huang, X. & Huang, Y. Cellsnp-lite: an efficient tool for genotyping single cells. Bioinformatics 37, 4569–4571 (2021).

114. Putri, G. H. et al. Extraction and quantification of lineage-tracing barcodes with NextClone and CloneDetective. Bioinformatics (2023).

115. Gayoso, A. et al. A Python library for probabilistic analysis of single-cell omics data. Nature Biotechnology 40, 163–166 (2022).

116. Subramanian, A. et al. Gene set enrichment analysis: a knowledge-based approach for interpreting genome-wide expression profiles. Proc. Natl. Acad. Sci. U. S. A. 102, 15545–15550 (2005).

117. Liberzon, A. et al. The Molecular Signatures Database (MSigDB) hallmark gene set collection. Cell Syst. 1, 417–425 (2015).

118. Faure, L., Soldatov, R., Kharchenko, P. V. & Adameyko, I. scFates: a scalable python package for advanced pseudotime and bifurcation analysis from single-cell data. Bioinformatics 39, btac746 (2023).

119. La Manno, G. et al. RNA velocity of single cells. Nature 560, 494–498 (2018).

120. Bergen, V., Lange, M., Peidli, S., Wolf, F. A. & Theis, F. J. Generalizing RNA velocity to transient cell states through dynamical modeling. Nat. Biotechnol. 38, 1408–1414 (2020).

121. Heumos, L. et al. Pertpy: an end-to-end framework for perturbation analysis. Bioinformatics (2024).

122. Howard, D. M. et al. Genome-wide meta-analysis of depression identifies 102 independent variants and highlights the importance of the prefrontal brain regions. Nat. Neurosci. 22, 343–352 (2019).

123. Trubetskoy, V. et al. Mapping genomic loci implicates genes and synaptic biology in schizophrenia. Nature 604, 502–508 (2022).

124. Mullins, N. et al. Genome-wide association study of more than 40,000 bipolar disorder cases provides new insights into the underlying biology. Nat. Genet. 53, 817–829 (2021).

125. Demontis, D. et al. Genome-wide analyses of ADHD identify 27 risk loci, refine the genetic architecture and implicate several cognitive domains. Nat. Genet. 55, 198–208 (2023).

126. Bellenguez, C. et al. New insights into the genetic etiology of Alzheimer’s disease and related dementias. Nat. Genet. 54, 412–436 (2022).

127. Cao, J. et al. The single-cell transcriptional landscape of mammalian organogenesis. Nature 566, 496–502 (2019).

